# Local adaptation to novel habitats and climatic changes in a wild tomato species via selective sweeps

**DOI:** 10.1101/2021.09.24.461657

**Authors:** Kai Wei, Gustavo A Silva-Arias, Aurélien Tellier

**Author notes:** these authors contributed equally (Kai Wei, Gustavo A Silva-Arias). Corresponding author: Gustavo A Silva-Arias.

## Abstract

- Positive selection is the driving force underpinning local adaptation, and leaves footprints of selective sweeps at the underlying major genes. Quantifying the timing of selection and revealing the genetic bases of adaptation in plants species occurring in steep and varying environmental gradients is crucial to predict a species’ ability colonize new niches.
- We use whole genome sequence data from six populations across three different habitats of the wild tomato species *Solanum chilense* to infer the past demographic history and search for genes under strong positive selection. We then correlate current and past climatic projections with the demographic history, allele frequencies, the age of selection events, and distribution shifts.
- We find evidence for several selective sweeps targeting regulatory networks involved in root hair development in low altitude, and response to photoperiod and vernalization in high altitude populations. These sweeps occur in a concerted fashion in a given regulatory gene network at particular periods of substantial climatic change.
- We decipher the genetic bases and the timing of local adaptation during plant colonization of semi-arid habitats using a unique combination of genome scans for selection and modelling of past climatic data.

## Introduction

Adaptation to abiotic conditions often occurs by means of positive selection. In heterogeneous environments, however, plants may be strongly influenced by locally variable selection. This can lead to divergence of populations at key loci (Savolainen *et al*., 2013; Tiffin & Ross-Ibarra, 2014), and results in trade-offs where native alleles show a fitness advantage relative to foreign alleles (antagonistic pleiotropy) culminating in local adaptation (Kawecki & Ebert, 2004). Positive selection underlies as well plant adaptation when colonizing new habitats (Savolainen *et al*., 2013; Tiffin & Ross-Ibarra, 2014), and/or when the environment changes in time at a given location (Polechová *et al*., 2009). With recent technological advances, it becomes possible to obtain genomes of many individuals across different populations to reveal the genetic bases underpinning adaptation to abiotic stress and/or gradient of stress. This can be achieved by genome scans for genes exhibiting signatures of (positive) selection in genome-wide polymorphism data, correlation between allele frequencies and environmental variables (e.g. RDA analysis), and/or genome-wide association studies with relevant phenotypes (review in *e*.*g*. (Savolainen *et al*., 2013; Josephs *et al*., 2017; Fagny & Austerlitz, 2021). Revealing the genetic bases of adaptation is not only important from an evolutionary biology perspective, but also to predict a species’ ability to colonize new niches as well as for applications to agriculture, whereby crops could be improved for stress tolerance using key adaptation genes found in related wild species.

Phenotypic traits of tolerance to abiotic stresses, such as drought, salt, cold, involve a set of complex and intertwined physiological, molecular, biochemical, and hormonal mechanisms and signals (Tardieu & Tuberosa, 2010), and therefore are complex (polygenic) traits encoded by many genes involving several gene networks or pathways. There has been a growing interest in the evolution of such polygenic traits recently, with several theoretical predictions emerging regarding the speed and genetic architecture of adaptation to either 1) the local optimum of a newly colonized habitat (Chevin *et al*., 2010), or 2) the moving environmental optimum, that is a changing environment in time at a given location (Polechová *et al*., 2009; Matuszewski *et al*., 2014; Jain & Stephan, 2017a). Under large enough population sizes and strong shift in the environmental optimum, both models predict that more significant steps of adaptation occur first at sites with strong selective coefficients, possibly generating selective sweeps (Chevin *et al*., 2010; Matuszewski *et al*., 2014; Jain & Stephan, 2017a). The so-called (hard) selective sweeps are polymorphism patterns (footprints) in the genome due to the rapid (tens to hundreds of generations) fixation of advantageous alleles and the associated hitchhiking effect (Smith & Haigh, 1974; Kim & Stephan, 2002). In other words, the theory of selective sweeps is not incompatible with that of polygenic selection (Barghi *et al*., 2020), and different number of major genes exhibiting selective sweep signatures are expected to underlie fast and strong adaptation of complex (polygenic) traits. The number and identity of these genes depends on the distribution of selection coefficients among the multiple genes involved in the traits, the efficiency of selection (a function of effective population size and recombination rate), the architecture of the traits, place of genes in gene networks/pathways, and gene pleiotropy (Jain & Stephan, 2017b; Barghi *et al*., 2020). Hard selective sweeps represent indeed one possible but more easily observable outcome of strong positive selection when considering that genes act in complex networks (polygenic quantitative traits) determining adaptation to new environmental conditions, for example abiotic stress. We focus here on detecting genes that have been under strong positive selection in the past and which underlie plant adaptation to new habitats or to changing environmental conditions in the wild relative tomato species *Solanum chilense*.

*Solanum chilense* is an outcrossing species found in southern Peru and northern Chile in mesic to very arid habitats (Nakazato *et al*., 2010). Its ancestral range is likely in marginal desert habitat of the coast and mid-altitude ‘pre-cordillera’ region (800 - 2000 m altitude) of southern Peru. *S. chilense* colonized independently two different southern, but arid, isolated regions around the Atacama desert at different time periods (Fig. 1a) (Böndel *et al*., 2015; Stam *et al*., 2019b): an early divergence (older than 50,000 years ago [ya]) with the colonization of coastal habitats (in Lomas formations), and a more recent lineage divergence (less than 25,000 ya) restricted to highland altitudes (above 2,400m) of the Andean plateau. Signatures of natural selection (positive or balancing) at genes involved in stress adaptation were found when scanning few candidate genes for biotic and abiotic stress response (Xia *et al*., 2010; Fischer *et al*., 2011; Böndel *et al*., 2015; Nosenko *et al*., 2016; Böndel *et al*., 2018; Stam *et al*., 2019b). In the present study, we obtain full genome sequence data for 30 diploid and highly heterozygous plants from six populations representing the three main habitats of the species (Fig. 1a, defined in Böndel et al. 2015): the central group (area of origin at low to high altitude, denoted as group C), south-coastal (SC) group, and south-highland (SH) group. The south-highland group strongly differs from the central group for its current climatic conditions (higher daily and annual temperature ranges, summer potential evapotranspiration and solar radiation), while the south-coastal appears as only marginally different from the environment prevailing in the central group (higher minimum temperature in summer and winter, and frequent fog episodes) (Fig. 1b). Our aims are first to infer accurately the past demographic history of the species colonization and to reconstruct recent dynamics of the species’ distribution range in response to climatic history. Second, we conduct genome scans for selective sweeps and assign functions and gene network topology to these candidate genes. Third, we link climatic and genetic data at candidate genes using a genotype-environment association analysis to highlight the relevance of key gene regulatory networks (pathways) for adaptation. We finally discuss the history of adaptation in *S. chilense* and future empirical studies needed to test our results.

**Figure 1.**
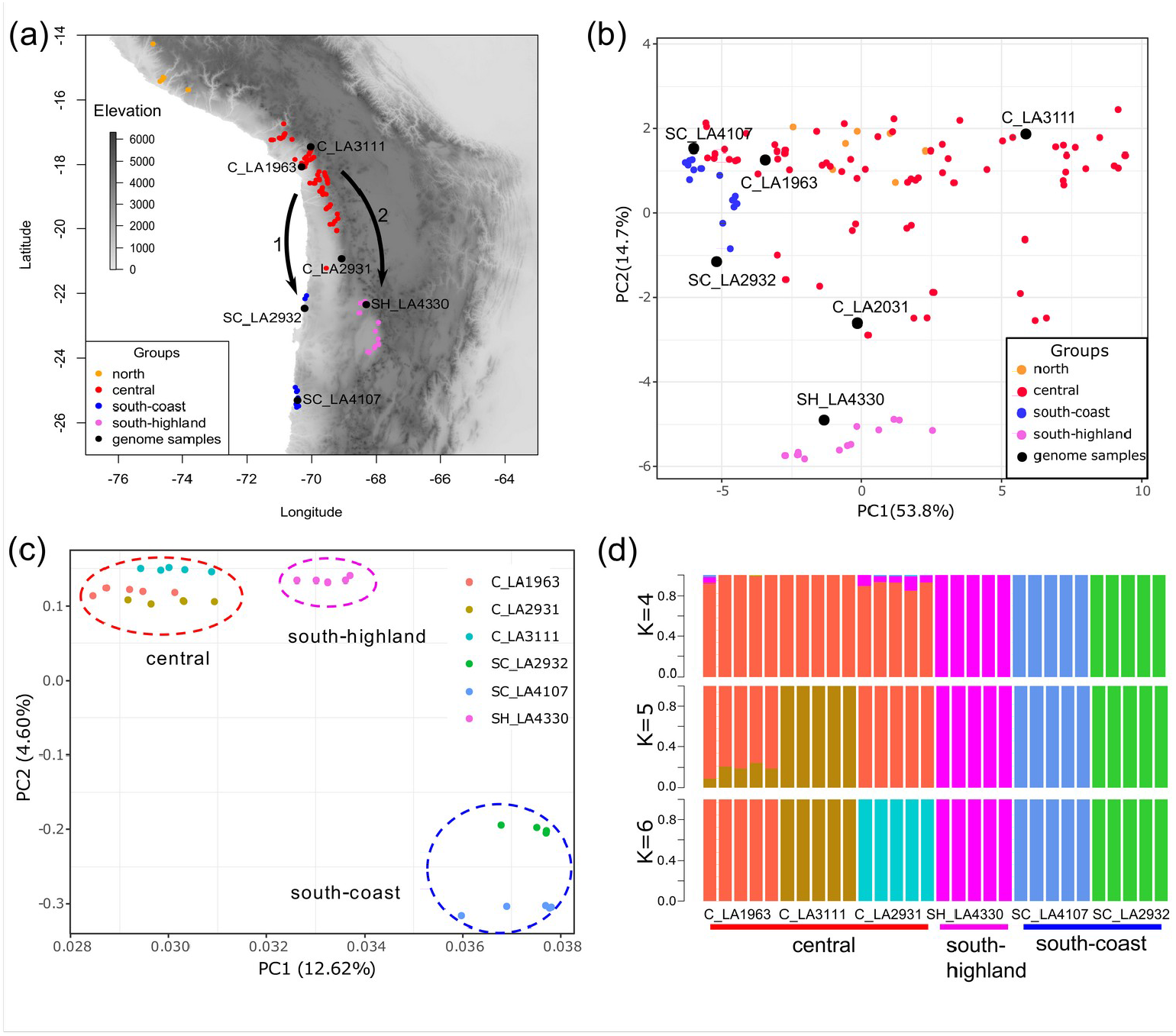
Geographic distribution and population structure of *Solanum chilense*. (a) Map with distribution of all *S. chilense* populations by the TGRC, the six *S. chilense* populations in this study (black circles), the four population groups (circles colours) and the two reconstructed southward colonization events, first to the south-coast and second to the south-highland (black arrows). (b) Principal components analysis of 63 current climatic variables from all *S. chilense* populations (Dataset S5). Population structure using SNP data based on (c) PCA and (d) Admixture analysis (optimal K value is 4; Fig S1b).

## Materials and Methods

### Sample collection, sequencing and bioinformatics

Plants were grown from individual seeds obtained from the Tomato Genetics Resource Center (TGRC, University of California, Davis, USA). We sampled five diploid plants from accessions C_LA1963, C_LA3111, C_LA2931, SH_LA4330, SC_LA2932, SC_LA4107 representing the three main geographic groups (C, SC, SH, Fig. 1a; Table S1). Plants were grown in standard glasshouse conditions. Genomic DNA was extracted using the DNA extraction kit from Quiagen and sequenced on an Illumina HiSeq 2500 with standard library size of 300 bp (Eurofins Genomics, Germany). The 30 *S. chilense* whole-genome sequencing data are available on ENA in BioProject PRJEB47577.

We performed quality control of the raw reads and trimmed calls with insufficient quality or adapter contamination. The clean reads were mapped to the *Solanum pennellii* reference genome (Bolger *et al*., 2014) available from Solanaceae Genomics Network using the Burrows-Wheeler Alignment tool (v0.7.16) with default settings (Li & Durbin, 2009) and sorted with Samtools (v1.5) (Wysoker *et al*., 2009). The raw alignments were then processed to add read groups, mark duplicates and fix mates. Variant calling was performed using the HaplotypeCaller tool of GATK (McKenna *et al*., 2010) (Auwera et al. 2013) with default parameters for each sample. Individual genomic variant files were then combined into a variant matrix with the GenotypeGVCFs tool and annotated based on the gene annotation of the *S. pennellii* reference (details in the Supplementary SI text 1).

### Population genetics analyses and inference of demographic history

For all population genetics analyses, we used *S. pennellii* population LA716 as outgroup. We built a maximum likelihood (ML) phylogenetic tree and performed a principal component analysis (PCA) and the inference of population structure with ADMIXTURE (Alexander *et al*., 2009). Population genetics statistics namely nucleotide diversity (π), Tajima’s D and *F*_ST_ for each population (or pairs of populations), were calculated with ANGSD v0.937 (Korneliussen *et al*., 2014) over 100 kb sliding non-overlapping windows. The linkage disequilibrium (LD) levels were calculated per population as the genotype correlation coefficient (r^2^) between two loci using VCFtools (Danecek *et al*., 2011) with a maximum distance of 1,000 kb.

The demographic inference was conducted using the Multiple Sequentially Markovian Coalescent method (MSMC2) with phased VCF files and 40 hidden states (Malaspinas *et al*., 2016). The cross-coalescence analysis was performed for each pairwise comparison of genomes between pairs of populations to estimate the population separation history and the migration rate with MSMC-IM (Wang *et al*., 2020). Phasing were generated with SHAPEIT v2 under the linkage disequilibrium mode (Delaneau *et al*., 2012). We assumed generation time of 5 years (uncertainty interval 3-7) and a mutation rate per generation of 1×10^−8^ (uncertainty interval 5.1×10^−9^ – 2.5×10^−8^, based on Roselius *et al*., 2005), accounting for ambiguity in these estimates.

### Modelling present and past species distribution

We reconstructed and visualized the environmental space occupied by *S. chilense* extracting the environmental conditions at the current occurrence points and summarize them by PCA (Fig. 1b) (Legendre & Legendre, 2012). The environmental data include 63 climatic layers obtained from three public databases WorldClim2 (Fick & Hijmans, 2017), ENVIREM (Title & Bemmels, 2018), and the Consultative Group on International Agricultural Research (CGIAR) (Trabucco & Zomer, 2018) (Dataset S5). The PCA was performed by the *prcomp* function in R (R Core Team, 2020).

We then performed an ensemble modelling framework (Araujo & New, 2007) using the BIOMOD package (Thuiller *et al*., 2009, 2014) in R, including all known localities covering the entire geographic range of *S. chilense* using eight modelling algorithms, five cross-validation replicates, and ten pseudo-absence sampling sets, therefore completing a total of 400 models. Consensus niche models were obtained using a TSS-weighted average method to account for the predictive power of each fitted model. Models with low predictive power (TSS < 0.7) were discarded. All fitted suitability models were then projected to infer the distribution of suitable habitats of *S. chilense* under current climatic conditions and during the Last Glacial Maximum (LGM; ∼21 Kya).

### Genome-wide selection scans and statistical power

We identified selective sweeps using biallelic SNPs by SweeD (Pavlidis *et al*., 2013) and OmegaPlus (Alachiotis *et al*., 2012). The CLR statistics in SweeD were calculated with default parameters with 10 kb intervals. OmegaPlus statistics (ω) were computed at 10 kb intervals. We specified a minimum window of 10 kb and a maximum window of 100 kb to be used for computing LD values between SNPs, respectively. Outlier CLR and ω statistics indicative of a selective sweep are defined by comparison to the genome-wide distribution values. To reduce false-positive outliers derived from demographic processes, the cut-off values of the CLR and ω statistics to determine outlier windows were defined by coalescent simulations of the inferred demographic history. The maximum value of each statistic was extracted from each simulated dataset, and we thus obtained a distribution of 10,000 maximum values for each statistic. The 95th percentile of this maximum distribution was specified as the thresholds to identify outlier windows. We used the coalescent simulator SCRM (Staab *et al*., 2015) to generate 10,000 neutral datasets of 10 Mb based on the demographic history of each population and assuming a varying recombination rate every 100 kb within each 10 Mb simulated block (recombination rate varied between 0.1*θ and 10*θ). Using the genomic coordinates, we then extracted only the overlap regions between the two methods, which are regarded as high confident selective sweep regions. As independent confirmation of the sweep regions, we used McSwan (Tournebize *et al*., 2019) to detect sweeps and estimate their age. McSwan was run with the same parameters as SweeD.

To evaluate the sensitivity of our sweeps detection pipeline, we simulated 1,000 selective sweeps assuming five *N*_e_ scaled selection coefficients from nearly neutral to strong selection (2*N*_e_*s*=0.1, 1, 10, 100 and 1000) for each of the six populations under the inferred demographic model, with five different sweep ages (8, 14, 29, 50, 71 thousand ya). We used the function *generate_pseudoobs* based on MSMS simulator implemented in the McSwan R package (Ewing & Hermisson, 2010; Tournebize *et al*., 2019). We then ran SweeD, OmegaPlus and McSwan on all simulated data sets using the same parameters and thresholds defined above to quantify the percentage of sweeps detected per population (and age of the detected sweeps).

### GO enrichment analysis and gene networks

Due to the lack of a complete gene function annotation database, we performed a BLASTX against the NCBI database of non-redundant proteins (nr) screened for green plants (e-value cutoff was 10^−6^) and used Blast2GO to assign GO terms for each gene identified in the genome scan analysis (Conesa *et al*., 2005; Conesa & Götz, 2008) as well as performed a blast to the *A. thaliana* dataset TAIR10 separately to remove redundant terms (Berardini *et al*., 2015). The false discovery rates (FDR) were calculated to estimate the extent to which genes were enriched in given GO categories (significance cutoff of P-values < 0.05). For each of the genes enriched in specific biological processes, we retrieved the interacting gene neighbours using GeneMANIA (Warde-Farley *et al*., 2010). We generated aggregate interaction networks in GeneMANIA, based on physical interactions, predicted and co-expression. Finally, we performed hierarchical clustering and manually optimized the weighted value cutoff for displaying the gene network.

### Redundancy analysis (RDA)

We tested for genotype-environment association (GEA) using RDA (Capblancq & Forester, 2021) using the *rda* function from the vegan package implemented in R (Oksanen *et al*., 2015), modelling genotypes as a function of the same climatic predictor variables used for the niche reconstruction analyses, and producing constrained axes and representative predictors. Multi-collinearity between representative predictors was assessed using the variance inflation factor (VIF) and since all predictor variables showed VIF < 20 none were excluded. This may still cause some collinearity, but it is beneficial to find more connections between genotypes and environments. The significance of RDA constrained axes was assessed using the *anova*.*cca* function and significant axes were then used to identify candidate loci (P < 0.001). Candidate loci were identified using 2.5 standard deviation as cut-off (two-tailed p-value = 0.012). In order to measure the rate of false-positive associations due to the demographic history, we also performed the same RDA analysis using 1) a set of 1,000 randomly chosen SNPs from non-sweep regions, and 2) polymorphism data from the neutral simulations used to calibrate the SweeD and OmegaPlus thresholds.

## Results

### Past colonization events and climatic variations in *Solanum chilense*

We sequence whole genomes of 30 heterozygous plants from *S. chilense* from six populations (C_LA3111, C_LA1963, C_LA2931, SC_LA2932, SC_LA4107, SH_LA4330) (Fig. 1a; Table S1) and used the reference genome assembly of *S. pennellii*. All 30 *S. chilense* individuals show high-quality sequence and mapping scores with more than 97% of mapping paired reads, individual genome coverage ranging between 16 to 24 reads per base, and >70% genome coverage per sample (Dataset S1). After SNP calling and stringent filtering, a total of 34,109,217 SNPs are identified across all 30 samples (Table S2) for a genome size estimated approximately to 914Mb (Stam *et al*., 2019a). Phylogenetic analysis, principal component analysis (PCA) and population genetics statistics (Fig. 1c, S1, S2; Table S3, S4) support the population structure into three genetic groups, confirming the results in (Böndel *et al*., 2015): a central group (C_LA1963, C_LA3111, C_LA2931), the south-highland group (SH_LA4330), and the south-coast group (SC_LA2932 and SC_LA4107). The two south-coast populations constitute independent groups (best K=4; Fig. 1d, S1b-d). Only the individuals of the population C_LA2931 (the southernmost of the central group) display small admixed ancestry coefficients (< 5%) with the south-highland group (SH_LA4330, Fig. 1d). There is no significant correlation between genetic (pairwise Nei’s distance) and geographical distance (Pearson test, r = 0.35, P = 0.20; Fig. S1e).

As we confirm that *S. chilense* independently colonized the coastal and highland southern habitats from a lowland area located north of the central group region (Böndel *et al*., 2015; Stam *et al*., 2019b), we further refine our estimates of the historical changes in effective population size (*N*_*e*_, Fig. 2a, S3), divergence and potential post-divergence gene flow (Fig. S4), and finally construct a consensus demographic model (Fig. 2b; Dataset S2; see Fig. S3 accounting for mutation rate and generation time uncertainties). These estimates are compared in Fig. 2b with the reconstructed past climatic variation highlighting five marine isotope stages (MIS) climatic periods (Lisiecki & Raymo, 2005; Ritter *et al*., 2019). The two south-coast populations found in Lomas habitats (SC_LA2932 and SC_LA4107) show early divergence consistent with the admixture analysis (during the Last Inter-Glacial period, MIS5) likely from the lowland area of the central group (C_LA1963). The colonization of the highland likely occurred later, first in the central group region (C_LA3111, C_LA2931) between the last interglacial and Last Glacial Maximum (LGM) periods (ca. 75-130 kya, MIS3-4) and then with further colonization of southern highlands (from 30 kya, MIS1-2, SH_LA4330). All populations show a moderate effective size reduction matching with the estimated time of the LGM characterized as a cold and dry period and supported by a contraction of the suitable habitats to a narrow strip in lower altitudes, and a subsequent expansion thereafter (Fig. 2a,c). Indeed, the local habitat at the current location of C_LA2931 and SH_LA4330 was likely unsuitable for the establishment of southern highland populations until 15 kya (after the LGM, *i*.*e*. during MIS1-2; Fig. 2c). The lower genetic diversity of the south populations (and estimated *Ne*) is thus due to a mild colonization bottleneck during the southward expansion (Fig. 2a,c, Fig. S2; Table S3). Both south-coast populations show consistent signals of long-term history of colonization, subsequent isolation with negligible gene flow, and possible local specialization to sparsely suitable Lomas habitats along the coast (Fig. 2b,c, S3).

**Figure 2.**
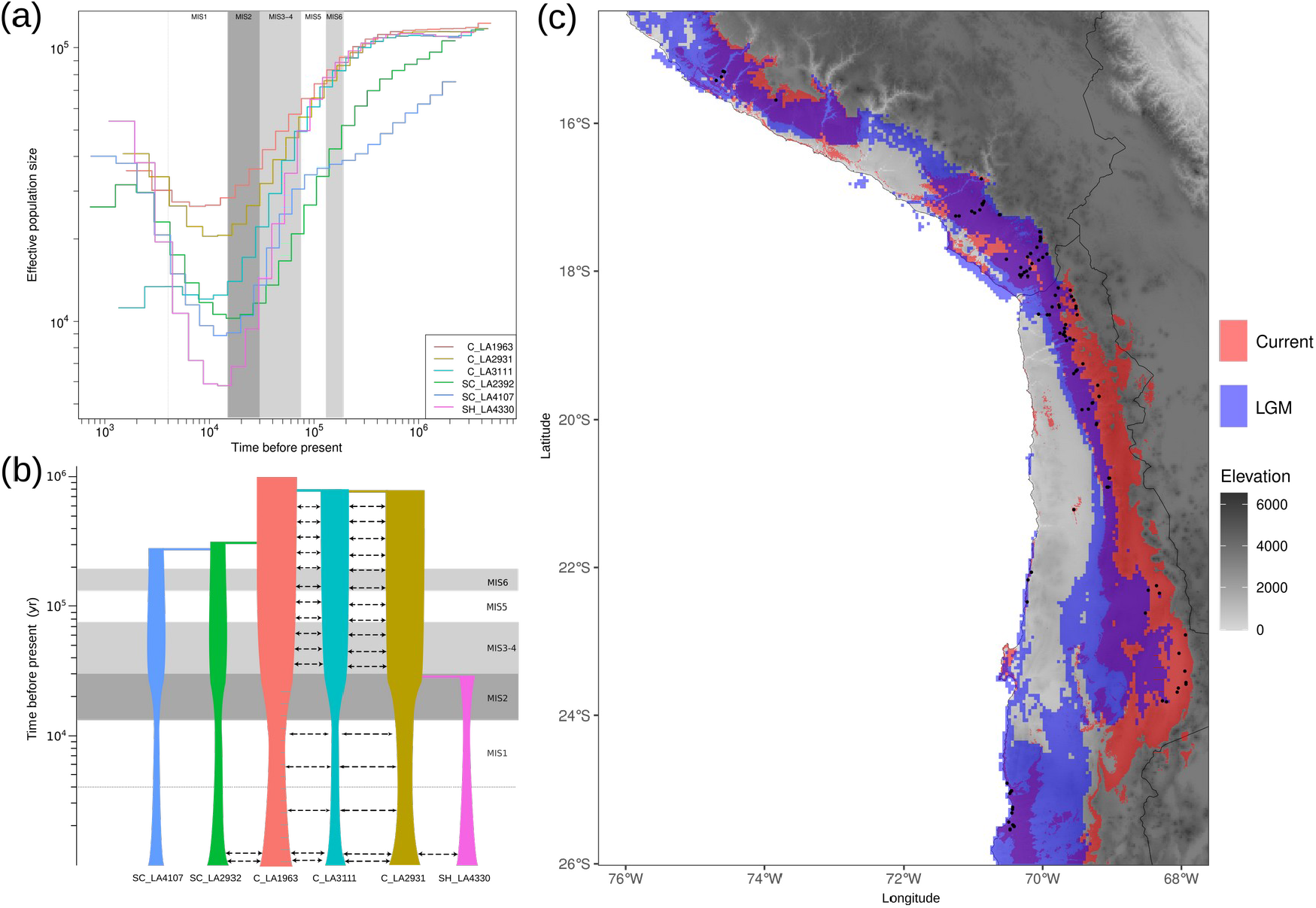
Demographic history and species distribution model of *S. chilense* for current and Last Glacial Maximum (LGM) climate conditions (a) The estimation of historical patterns of effective population size (*N*_*e*_) for 10 pairwise genome comparisons per population using the MSMC model. (b) Interpreted demographic scenario for the six samples populations of *S. chilense* including likely estimations of effective population size, divergence times and gene-flow. The width of the boxes represents the relative effective population size, arrows represent the migration between population pairs. Grey background boxes indicate five Marine isotope stages (MIS) climatic periods. (c) Overlay of the reconstruction of the distribution model for *S. chilense* using current climatic variables (red) and LGM past climatic variables (blue). Darker color of the gradient indicates higher suitable habitat for a given climatic period.

The divergence between the central group populations (during MIS3-4) occurs in parallel to the colonization of the coastal habitat (Fig. 2b, S4), but before the colonization of the south-highland (SH_LA4330). Moreover, strong post-divergence gene-flow and low differentiation are found in the central group, especially among the pairs C_LA1963-C_LA3111 and C_LA3111-C_LA2931 (Fig. S2c, S4), consistent with their geographical and/or environmental proximity (Fig. 1a,b) and the range contraction during the LGM (MIS2 in Fig. 2). The colonization of high-altitude regions in the central group is thus accompanied by high levels of gene flow despite these populations ranging across a large altitudinal gradient (2500m of altitude difference between C_LA1963 and C_LA3111 or C_LA2931). The divergence history results in the south-coast and south-highland populations to be fairly isolated from one another (as separated by the Atacama desert) leading to the suggestion of an incipient speciation process (Fig. 2b, S4) (Raduski & Igić, 2021). In contrast to the study of Böndel *et al*., (2015), our smaller number of populations and the independent divergence histories of the two southern groups, does not allow us to find a significant signature of isolation by distance.

### Selective sweeps underpin local adaptation

In total, we find 2,921 candidate sweep regions with SweeD (mean size 212,858 bp +/-3,938) and 13,106 with OmegaPlus (mean size 59,618 bp +/-521) across all six populations (Table 1), yielding a total of 520 overlapping regions (mean size 41,082 bp +/-1,618). Although we calculate SweeD and OmegaPlus statistics by 10kb interval, we found in fact that the estimated sweeps in SweeD are larger than Omegaplus. Therefore, in most cases sweep regions identified from SweeD overlap with multiple sweep regions identified from OmegaPlus. These regions contain 799 protein-coding candidate genes assumed to be under positive selection (Fig. S5; Dataset S3). In SC_LA4107, we find 61 candidate genes and about 100 candidate genes are detected in each of the other four populations (Table 1). The largest number of candidate genes (354) is found in SH_LA4330 (Table 1), likely because the population has been established recently (Fig. 2a,b), and its habitat is ostensible different from the rest of the species range (Fig. 1b).

**Table 1.** The summary of genome scans and estimation of sweep age.

| Population | Genome Scans |  |  |  | Sweeps age |  |  |  |
| --- | --- | --- | --- | --- | --- | --- | --- | --- |
| | $N_{\text{SweeD}}$ | $N_{\text{OmegaPlus}}$ | $N_{\text{overlaps1}}$ | $N_{\text{genes1}}$ | $N_{\text{McSwan}}$ | $N_{\text{overlaps2}}$ | $N_{\text{genes2}}$ | $Age_{\text{mean}}$ (kyr) |
| C_LA1963 | 385 | 2,474 | 98<sup>b</sup> | 86 | 267 | 16 | 14 | 38 ± 16 |
| C_LA2931 | 517 | 2,268 | 109 | 125 | 355 | 24 | 28 | 20 ± 10 |
| SC_LA2932 | 374 | 1,717 | 46 | 101 | 302 | 15 | 29 | 36 ± 15 |
| C_LA3111 | 663 | 2,307 | 105 | 107 | 377 | 22 | 22 | 23 ± 11 |
| SC_LA4107 | 203 | 2,047 | 37 | 61 | 194 | 11 | 13 | 34 ± 10 |
| SH_LA4330 | 779 | 2,293 | 125 | 354 | 438 | 36 | 71 | 17 ± 8 |

$N_{\text{SweeD}}$ , number of outlier regions from SweeD; $N_{\text{OmegaPlus}}$ , number of outlier regions from OmegaPlus; $N_{\text{overlaps1}}$ , number of overlapping regions between SweeD and OmegaPlus; $N_{\text{genes1}}$ , number of candidate genes in overlaps1, and all candidate genes show in Dataset S3; $N_{\text{McSwan}}$ , number of outlier regions from McSwan; $N_{\text{overlaps2}}$ , number of overlapping regions between McSwan and overlaps1; $N_{\text{genes2}}$ , number of genes in overlaps2; $Age_{\text{mean}}$ , mean age ± standard deviation of overlaps2.

We present here two arguments supporting the fact that our cutoff values are well designed on the basis of the population demography to reasonably discriminate between demography and selection signals (as shown in Huber *et al*., 2014). First, comparison of genome-wide genetic diversity statistics (π and Tajima’s D) between the observed data and the neutral simulations show that our demographic scenario captures well the genomic diversity patterns in all populations (Fig. S2). Second, we estimate by simulations the accuracy (statistical power) to detect sweeps under our demographic model and a range of selection coefficients and sweep ages. The accuracy is found to be between 0 - 9.3% for nearly neutral to weak selection coefficients (2 *N*_*e*_ s = 0.1 - 1), 1.1 - 90.5% for strong selection (2 *N*_*e*_ s = 10 - 100), and 64.7 - 93.8% for very strong selection (2 *N*_*e*_ s = 1000) across populations and for each method (Figure S6). SH_LA4330 exhibits even relatively high statistical power compared to the other populations (Figure S6). Furthermore, the detection power increases for intermediate sweep ages (14-50 thousand ya; Figure S6). This demonstrates that our defined thresholds for sweep detection are conservative and allow minimizing the rate of false positives, at the small cost of not detecting all selective sweeps, especially if the selection coefficients are too small and the sweep are too recent or too old (Figure S6). Further, only a few candidate genes are shared among different populations, with the central and south-highland populations sharing a small number of candidate genes, while almost none are shared between the two south-coastal populations (Fig. S5c). This lack of common candidate genes among populations is likely due to 1) the high effective population sizes (Fig. 2a) generating new variants across many genes which are then differentially picked up by selection across different populations, 2) the relatively old inter-population divergence and timing of local adaptation, and 3) the marked environmental differences between the central and the two southern regions promoting sweeps in different pathways.

An overview of population genetics statistics shows that our candidate regions exhibit typical characteristics (lower nucleotide diversity, higher LD, more negative Tajima’s D and higher pairwise *F*_*ST*_ values) of positively selected regions when compared to the genome-wide statistics (see SI Text 2; Fig. S6; Table S3, S4). Furthermore, we find an overlap between our candidate genes under selective sweep and genes exhibiting signals of positive selection in previous studies in *S. chilense* which are based on few chosen genes, different plants, different populations and different sample sizes than ours. Among our candidate genes, we indeed find three genes (*JERF*3, *TPP* and *CT*189) involved in abiotic stress tolerance such as salt, drought or cold (Böndel *et al*., 2015) as well as three NLRs (nucleotide binding leucine rich repeat, SOLCI006592800, SOLCI001535800, SOLCI005342400) possibly linked to resistance to pathogens (Stam *et al*., 2019b). We also find that two of the seven most up-regulated genes under cold conditions in a transcriptomic study of *S. chilense* (Nosenko *et al*., 2016) do appear in our selection scan in high altitude populations: *CBF*3 (Solyc03g026270) in C_LA2931, and *CBF*1 (Solyc03g026280) in SH_LA4330. These results indicate that our genome-wide selective sweep scan generalizes the previous selection studies in *S. chilense* and supports the functional relevance of our candidate genes.

### Gene regulatory networks underlying local adaptation in *S. chilense*

A Gene Ontology (GO) enrichment analysis of the 799 candidate genes reveals common GO categories in all populations for basic cell metabolism, immune response, specific organ development, and response to external stimuli (Fig. S7). Most interesting, are four GO categories restricted to populations with distinct habitats: (i) root hair cell differentiation functions are enriched in 15 candidate genes, only in the three coastal populations (C_LA1963, SC_LA2932 and SC_LA4107); (ii) response to circadian rhythm, photoperiodicity and flowering time are enriched in 12 candidate genes in two high-altitude (C_LA3111 and SH_LA4330) and a south-coast (SC_LA2932) populations; (iii) vernalization response is enriched in eight candidate genes in the three high-altitude populations (C_LA2931, C_LA3111, SH_LA4330), and (iv) protein lipidation is enriched in seven candidate genes in the south-highland population (SH_LA4330). Based on the wealth of available data in cultivated tomato, *S. pennellii* and *A. thaliana*, we further study the gene regulatory networks to which the candidate genes belong.

For adaptation to high-altitude conditions, 15 candidate genes are interconnected in a flowering gene network, which is itself subdivided into two sub-networks related to flowering, photoperiod and vernalization control pathways (Fig. 3a; Dataset S4). Photoperiod responsive genes can sense changes in sunlight and affect the circadian rhythm to regulate plant flowering (Johansson & Staiger, 2015; Song *et al*., 2015), while vernalization genes regulate the flowering and germination through long-term low temperature (Guo *et al*., 2018; Xu & Chong, 2018; Iida & Mähönen, 2020). These two sub-networks are connected through several key genes, some of which appear as candidate genes entailing local adaptation in our populations: FL FLOWERING LOCUS C (FLC or AGL25), FLOWERING LOCUS T (FT) and AGAMOUS-LIKE genes (AGL) (Fig. 3a,b). These key genes are essential regulators acting on the flowering regulation pathway (Michaels & Amasino, 1999; Sheldon *et al*., 2000; Turck *et al*., 2008; Putterill & Varkonyi-Gasic, 2016). Remarkably, some candidate genes in the recently diverged south-highland population (SH_LA4330) aggregate into an independent network involved in circadian rhythm regulation, connected to the photoperiod network by JUMONJI DOMAIN CONTAINING 5 (JMJD5) also a candidate gene in C_LA3111 (Fig. 3a). In the central-highland population (C_LA3111), several other candidate genes of the photoperiod network also regulate circadian rhythm and flowering time. The three high-altitude populations (C_LA3111, C_LA2931, and SH_LA4330) have candidate genes of the AGAMOUS-LIKE (AGL) gene family in the vernalization network (Fig. 3a). We also note that the network of protein lipidation genes appears to be related to the synthesis of fatty acids in the south-highland population (Fig. 3d; Dataset S4). We speculate here that this latter adaptation may be related to adaptation to lowest-temperature stress of SH_LA4330 (Dataset S5) (Maksimov *et al*., 2017; Jiang *et al*., 2018). Adaptation to high altitude involves the regulation of the flowering, including photoperiod and vernalization pathways, but through different genes in different populations, while cold stress and its consequence (adaptation in lipidation pathway) may be relevant for adaptation to the highest altitudes (SH_LA4330).

**Figure 3.**
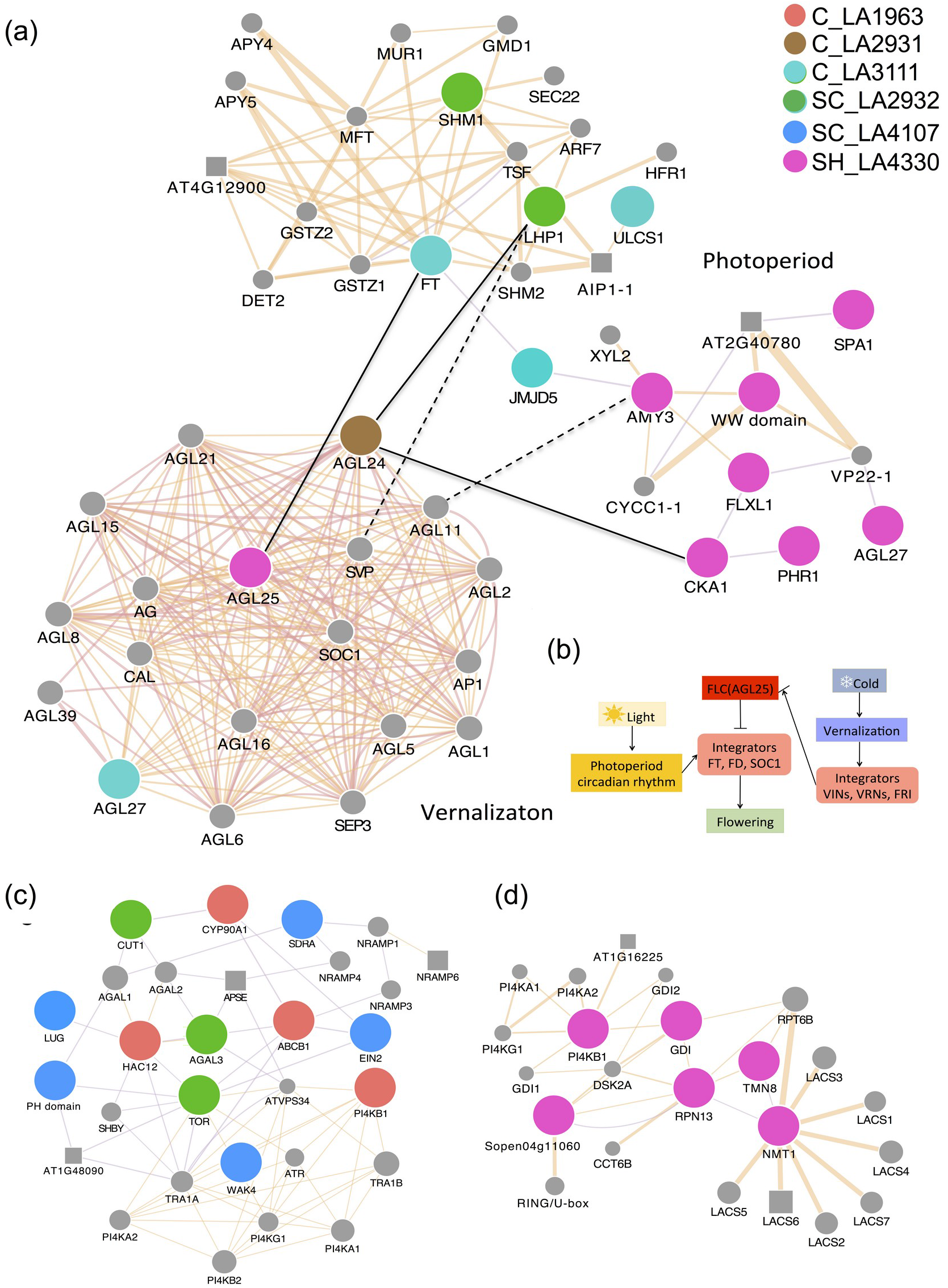
Interaction genetic networks of candidate genes. (a) The network of flowering regulation involved two sub-networks, photoperiod and vernalization pathways, for regulation of flowering. (b) The schematic diagram of flowering regulation involved photoperiod and vernalization is adapted from (Xu & Chong, 2018). “┤”indicates repressive effects on gene expression; “→”indicates promotive effects on gene expression. (c) The network of root development and cell homeostasis. (d) The networks of protein lipidation. Connections represent gene interactions based on physical interactions, informatics predictions and co-expression analyses. Connection thickness is proportional to weighted value of the connected genes. The black lines connected two sub-networks, genes under selection were connected by solid lines and other genes were connected by dashed line. Node colors correspond to genes were detected the different populations in genome scans. Gray circles, not detected in genome scan, but present in *S. chilense*; gray squares: not present in *S. chilense*.

Regarding adaptation to coastal conditions, we find 11 candidate genes related to root development and cellular homeostasis functions clustered in a single network (Fig. 3c; Dataset S4). We speculate that the drought and water shortage typical of the coastal conditions (Dataset S5) would promote the differentiation and extension of plant roots (Xiong *et al*., 2013; Li *et al*., 2017). The cell WALL ASSOCIATED LINASE 4 (WAK4), a candidate gene identified SC_LA4107, acts as a linker of signal from the cell wall to the plasma membrane and thus serve a vital role in lateral root development (He *et al*., 1999; Lally *et al*., 2001). In addition to root development, we also find genes involved in cell homeostasis (Fig. 3c; Dataset S4), which would be critical for the coastal drought and salinity conditions to maintain the stability of the intracellular environment in the coastal habitats (Forni *et al*., 2017; Zhao *et al*., 2020).

### Candidate genes show genotype-environment associations

Our candidate loci are hypothesized to be responsible for adaptation to local climatic conditions, so we test for genotype-environment association (GEA) using redundancy analysis (RDA). We perform first a “present day” RDA using 144,713 SNPs from all candidate regions and 63 climatic variables representing current (present) conditions for temperature, precipitation, solar radiation, and wind (Dataset S5). We find that the two-first RDA axes are significant (ANOVA’s P < 0.001) and retain most (38% and 21%) of the putative adaptive genetic variance identified in the genome scans in all populations (Fig. 4a). Tables S6 and S7 summarize outlier SNPs in different RDA models and their correlation with climatic variables. In concordance with the PCAs of both climatic and genomic variation (Fig. 1b,c), the two main RDA axes cluster the individuals into three groups corresponding to the main geographical regions (central, south-highland, and south-coastal) supporting that those axes synthesize the principal selective pressures for local spatial adaptation along with the species distribution (Fig. 4a; Table S7). RDA1 represents the differentiation of the two south-coast populations in correlation with higher precipitation of the coldest quarter (Bio19) and annual variation of solar radiation (CV_R) and RDA2 summarizes a climatic gradient differentiating the south-highland population mainly driven by annual potential evapotranspiration (annualPET) and temperature annual range (Bio7) (Table S7).

**Figure 4.**
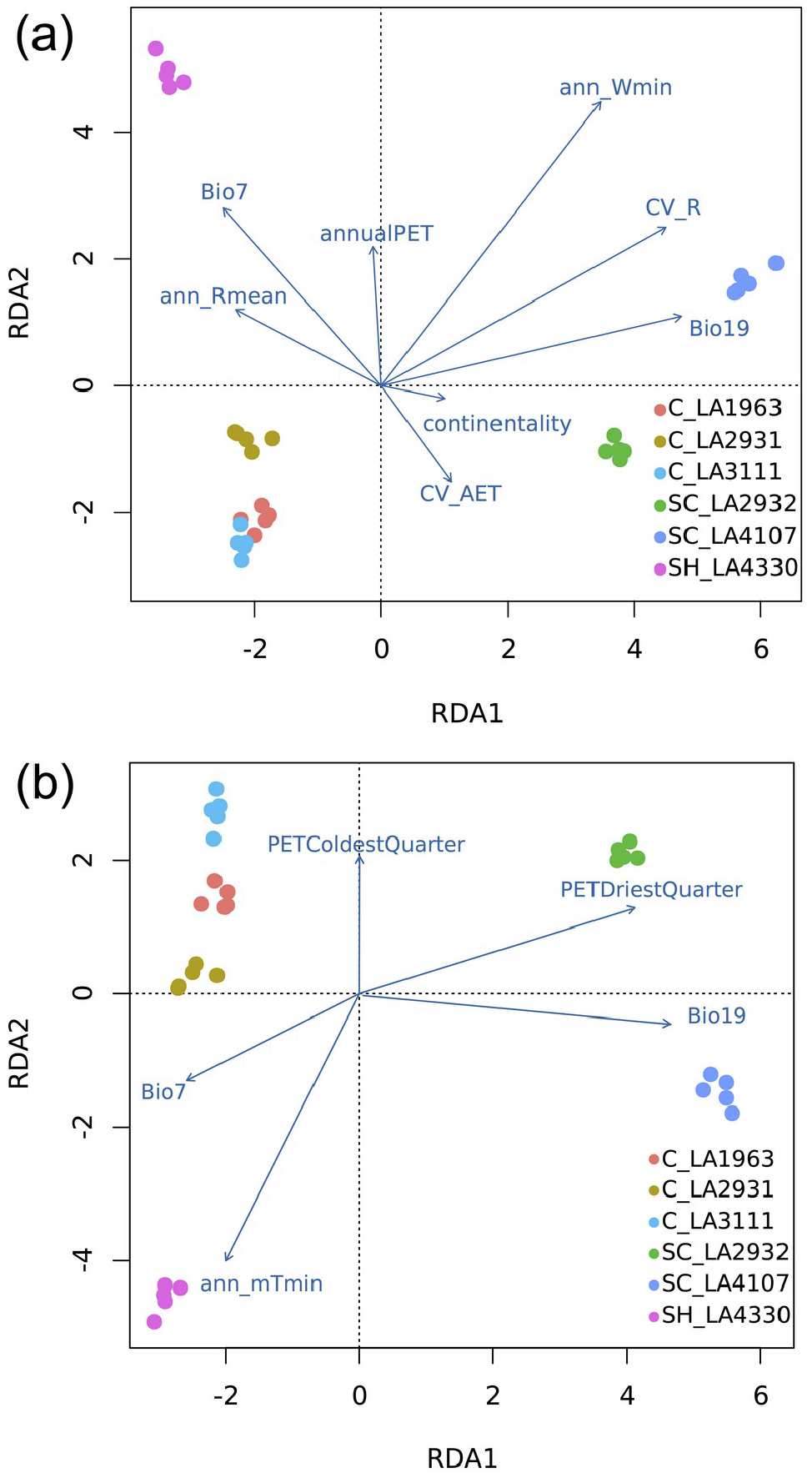
Redundancy analysis (RDA) ordination bi-plots between the climatic variables, populations and the genetic variants in all candidate sweeps. (a) RDA using current climatic variables. (b) RDA using LGM climatic variables. Arrows indicated the direction and magnitude of variables correlated with the populations. The abbreviations of climatic variables are shown in Dataset S5.

Further RDA analyses based on gene variants of the GO categories circadian rhythm-photoperiodism, vernalization, root-hair differentiation, and protein lipidation highlight combinations of climatic variables and genetic variants related to local spatial adaptation (Fig. S8a,c,e,g). These analyses show that the two main RDA axes explain 40% of the variation. Climatic variables representing temperature variability through the year such as temperature seasonality (Bio4) and temperature annual range (Bio7) are consistently correlated with adaptive variation of the south-highland population (Fig. 4a, S8). A total of 68 SNPs within candidate genes of the population SH_LA4330 are strongly associated with these two variables (Bio4, Bio7) in three of the RDA based on the GO categories (circadian rhythm-photoperiodism, vernalization, and protein lipidation; Dataset S6). The RDA based on the root-hair differentiation GO category exhibits a strong differentiation between lowland and highland populations based on atmosphere water vapour availability variables (ann_Vmin, ann_AET; Fig. S8e). Note that the RDA testing for false positives implemented on 1,000 random SNPs from non-sweep regions and neutral simulations produced no significant RDA axes and correlations with any climatic variable.

To assess the occurrence of selection in *S. chilense* as a response to past climatic changes, we implement an “LGM” RDA using 37 climate variables projected to the Last Glacial Maximum conditions (Fig. 4b, S8b,d,f,h; Dataset S5). This LGM RDA analysis aims to uncover additional genomic variation selected in response to temporal climatic changes and underlying the niche expansion towards the south-highland region (Fig. 1b). The LGM RDA analyses capture a smaller proportion of the genetic variability in the first two constrained axes (30%) compared to that using the current climatic variables. About 30% outlier SNPs are identified in genomic regions correlated with past climatic variables and not with the current variables (Table S6, S7). For example, the central populations C_LA3111 and C_LA2931 are separated in the past RDA of vernalization genes using LGM climatic variables indicating that warmer climate after LGM may drive gene flow among central populations as seen in the current RDA (Fig. 2b, S4, S8c,d). The past RDA of LGM climatic variables unveils that high-altitude populations, especially SH_LA4330, have SNPs correlating with temperature (*i*.*e*. annual mean minimum temperature; ann_mTmin, and temperature annual range; Bio7) whereas coastal populations SNPs do correlate with precipitation and potential evapotranspiration of the coldest and driest seasons (Fig. 4b). We advise caution in interpreting the LGM RDA results as these are based on climatic values from locations that likely had little or no population occurrence in the past, especially those in highland areas (but only mild bottlenecks could suggest local persistence, Fig. 2a,c). We suggest that this analysis is nevertheless useful for identifying alleles that arose in response to sudden changes in adaptive climatic optima during glacial-interglacial transitions, especially in the highland populations.

### Age of selective sweeps and timing of selection

We finally estimate the age of 112 selective sweep regions, that is the time since the fixation of the selected alleles, that overlap between the three positive selection detection methods (McSwan, SweeD and OmegaPlus, Table 1). These regions contain 175 genes and exhibit a mean sweep age of ca. 28,000 years. The ages of sweeps range from as early as 65 kya up to 2.5 kya (Table 1, Fig. 5). The highland populations exhibit more recent sweeps (2.5 - 35 kya) than those at the coastal populations, consistent with the recent (re)colonization of higher altitudes (Fig. 5). The south-coastal populations exhibit older and large distributions of sweep age consistent with older events of colonization (2.5 - 65 kya). Regarding the key gene networks of relevance for local adaptation highlighted above (root hair, protein lipidation, vernalization and photoperiod), each of them exhibits a narrow range of sweep age values across several populations (Fig. 5). The averages of sweep ages observed (Table 1) are perfectly in line with the estimates obtained from the sweep simulations under our demographic model (Table S5), demonstrating that our statistical power is adequate to estimate sweep ages under the demographic model and that old sweeps in the highland populations cannot be recovered (even if they occurred) in contrast to the coastal populations.

**Figure 5.**
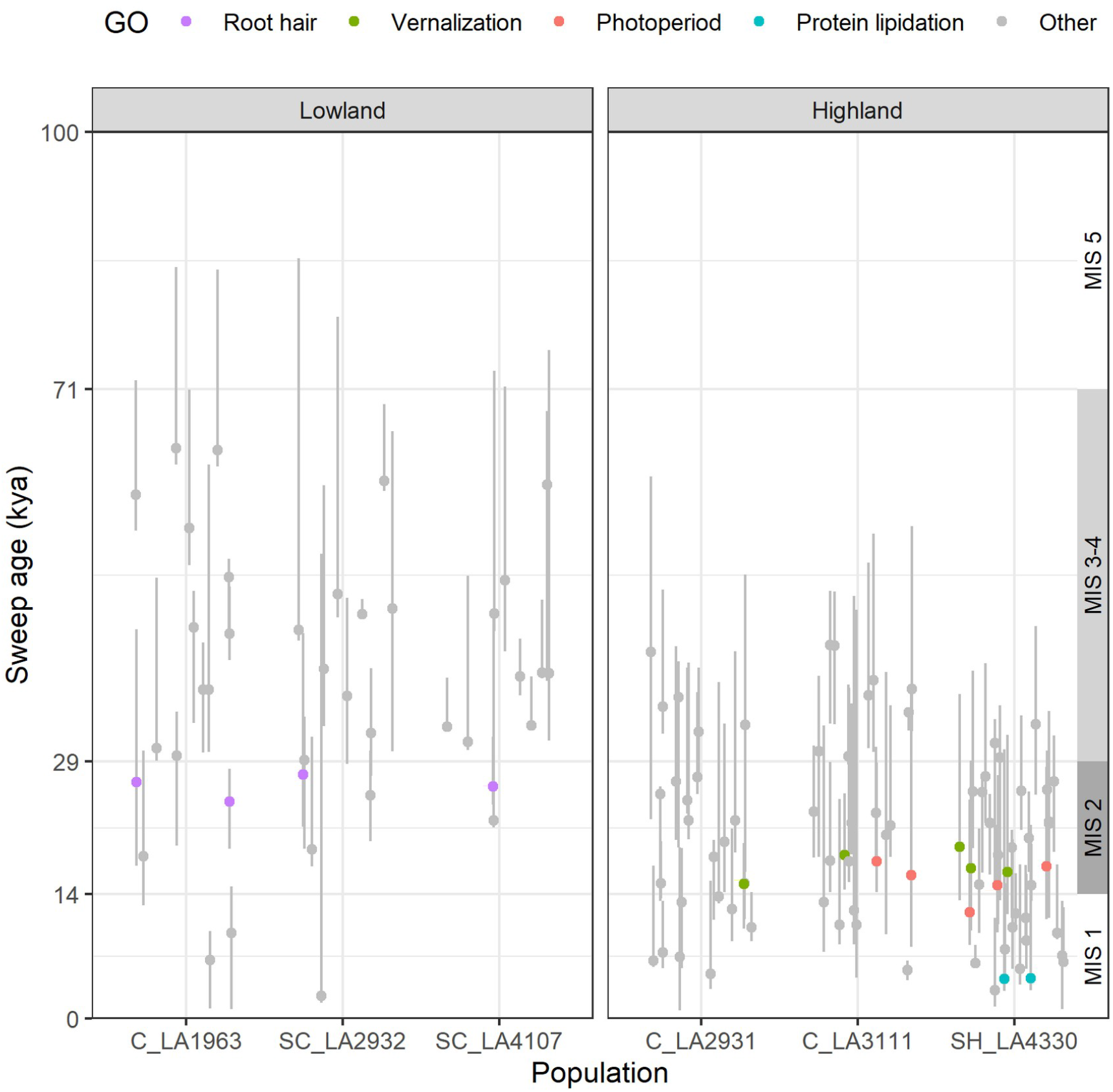
Distribution of estimated age of 112 selective sweeps highlighting five marine isotope stages (MIS) periods of climatic variation and sweeps containing genes within the four Gene Ontology categories related with local adaptation in *Solanum chilense*. The points represent mean age and lines the 95% confidence intervals. Generation time=5; µ=10^−8^.

## Discussion

Our study is the first to attempt to dissect in plants the complex selective processes and their genetic bases involved during and after the colonization of new highly stressful hyper-arid environments around the Atacama desert.

Taking our demographic and selection results altogether, we formulate the following scenario for the highland colonization. During the past colder climate phases (LGM-MIS2 at 30-15kya), the suitable areas of the species likely decreased at high altitude (Fig. 2c). We speculate that the populations were already established at high altitude before the LGM (MIS 3-5; Fig. 2) likely in the northern part of the range (from the location of C_LA3111 up to that of C_LA2931), before a contraction of the species range occurred towards lower altitudes during the LGM, and the subsequent colonization of new southern locations concluded ∼15kya (post-LGM, SH_LA4330). The highland populations (C_LA3111, C_LA2931, SH_LA4330) show adaptation by a burst, *i*.*e*. over a short time, of numerous selective sweeps across several gene networks (Fig. 5) but also show older selective events pre-dating the LGM (MIS3-4). Interestingly, the population SH_LA4330 exhibits selective sweeps in the vernalization and photoperiod which pre-date its establishment. These selective events likely occurred in the northern part of the range (C_LA3111, C_LA2931) during MIS2-4 acting as pre-adaptation requisite for colonizing the more divergent and extreme environments of the south-highlands (SH_LA4330; Fig. 1b).

The *S. chilense* lineage likely originates from coastal up to ‘pre-cordillera’ (800 - 2000 m altitude) habitats in southern Peru, explaining the early divergence and southward colonization process, accompanied by habitat fragmentation and contraction, which yields two highly isolated populations on the coast (Fig. 2b,c; SC_LA2932 and SC_LA 4107). The coastal colonization process seems to involve fewer sweeps than the adaptation to higher altitudes, for example a burst of selective sweeps in genes related to root anatomical traits during the LGM-MIS2 period (Fig. 5). We speculate here that these sweeps are due to temporal adaptation to changes in the habitat after colonization. However, some of the adaptive genomic signals in the coastal populations could be blurred due to the older divergence time and stronger drift (due to habitat fragmentation along the coast), or be incomplete/partial/soft sweeps (with small selection coefficients) which we do not detect (*e*.*g*. Garud *et al*., 2021).

We find between 60 and 350 selective sweeps per population, but contrary to our naïve expectations and previous findings in the literature, sweeps show a large distribution of ages, especially in the south-coastal populations. We suggest that several sweeps do occur concomitantly in a given gene pathway/network at a given time period, either to promote adaptation to a new habitat or in response to a moving environmental optimum (our climatic periods, temporal adaptation) as predicted under the polygenic model of adaptation (Polechová *et al*., 2009; Chevin *et al*., 2010; Matuszewski *et al*., 2014; Jain & Stephan, 2017a). Selective sweeps at genes with large selection coefficients can be observed because the populations of *S. chilense* exhibit large effective sizes (Fig. 2; Böndel *et al*., 2015), especially when compared to the small above ground abundance (census size) reported in these semi-arid habitats (Tellier *et al*., 2011). *S. chilense* is outcrossing and exhibits persistent seed banking. Both factors contribute to generate large effective population sizes by 1) decreasing linkage disequilibrium and the effect of linked selection, 2) buffering the negative impact of colonization bottlenecks, and 3) enhance the recovery post-colonization (Fig. 2; Tellier *et al*., 2011; Živković & Tellier, 2018). Therefore, the detection of old sweeps dating up to 65 kya for the coastal populations and up to 35 kya for the highland populations (pre-dating the recent post-LGM colonization; Fig. 2) is made possible and stretching beyond the theoretical limit of 0.1*N*_*e*_ computed without the effect of seed banking (Kim & Stephan, 2002).

As a word of caution, we focus on four main GO categories, which can be reliably associated with physiological traits likely underlying adaptation: root hair differentiation, vernalization, photoperiod, and protein lipidation. Pinpointing the regulatory or non-coding SNPs under selection was not possible with our sample sizes and functional information on many candidate genes in Figure 5 and S6 is still lacking to provide a complete picture. We indeed should not assume that all genes in the outlier windows are under selection, and therefore we designed a strategy in several steps to reduce the amount of potentially hitch-hiking genes. First, we reduce the set of candidate genes to only those in the overlapping regions of the outlier windows identified with different methods (SweeD and OmegaPlus, which rely on different summary statistics). Second, this subset was then reduced to set of genes that enriched biological functions showing physiological meaning based on the ecology of the populations (albeit avoiding the caveat described in Pavlidis *et al*., 2012). Third, we use the genotype-environment association analysis to focus only on a subset of outlier genes for demography and which correlate with key current and past climatic variables. We verified that the variants in the selected genes show the expected distributions (hallmarks of selective sweeps) in population genetics statistics compared to genome wide patterns. We acknowledge the limitations of genomic scans for selection in non-model species for which a recombination map is lacking and small sample size limit our ability to zoom in the sweep regions. Therefore, it is likely that our approach despite being conservative may have generated some false positives and missed some genes under selection. Furthermore, we focus here on selective sweeps resulting from strong positive selection as we cannot assess in our data the occurrence of weaker positive (polygenic) selection or signatures of soft or incomplete sweeps which would be favoured by the presence of seed banking (Živković & Tellier, 2018). Yet, we are confident that our candidate genes under selection are functionally relevant, as demonstrated by the overlap with previous studies (Böndel et al., 2015; Nosenko et al., 2016; Stam et al., 2019b).

We note also the possible bias in our results due to the use of accessions maintained and multiplied at TGRC (UC Davis, USA). Indeed accession multiplication in the glasshouse may change allele frequencies (SFS) and bias some of our demographic and selection inference. We provide in Figure S10 a summary of the previous data from (Böndel *et al*., 2015) containing the accessions of this study, in which we find that the maintenance at TGRC does reduce the number of rare alleles (and thus Tajima’s D), but only for accessions multiplied more than twice. As the accessions used here have been multiplied only once or twice, we consider that the bias may likely be minor on our inference. Nevertheless, our selection scans are not exhaustive and future work requires larger sample sizes as well as original material from *S. chilense* populations from the field to reveal the extent of positive and balancing selection in this species. To demonstrate whether the genes under positive selection contribute to local adaptation, further experimental work *in situ* and in common garden is needed.

## Supplementary information

Available as a single pdf with additional figures, tables and fully detailed methods.

## Acknowledgments

We thank three anonymous reviewers for their comments which helped to improve the manuscript. AT acknowledges funding from DFG (Deutscheforschungsgemeinschaft) Grant Number: 317616126 (TE809/7-1); KW was funded by the Chinese Scholarship Council, and GSA funded by the Technical University of Munich. We thank the Tomato Genetics Resource Center (TGRC) of the University of California, Davis for generously providing us with the seeds of the accession included in this study. We thank Daniela Scheikl for assistance with plant work and Christine Würmser for the Illumina sequencing.

## Author contributions

Conceptualization, GSA and AT; Software, Formal analysis and Investigation, KW and GSA, Writing KW, GSA and AT; Funding Acquisition, AT; Supervision, GSA and AT.

## Data availability

The raw pair-end sequencing genomic data used in all the analyses can be accessed at the European Nucleotide Archive (ENA) project accession PRJEB47577.

All code used in this study and other previously published genomic data is available at the sources referenced. The developed scripts and extra data formats are found on our GitLab repository: https://gitlab.lrz.de/population_genetics/wild-tomato-genomics.

