## Supplementary Information for "Local adaptation to novel habitats and climatic changes in a wild tomato species via selective sweeps"

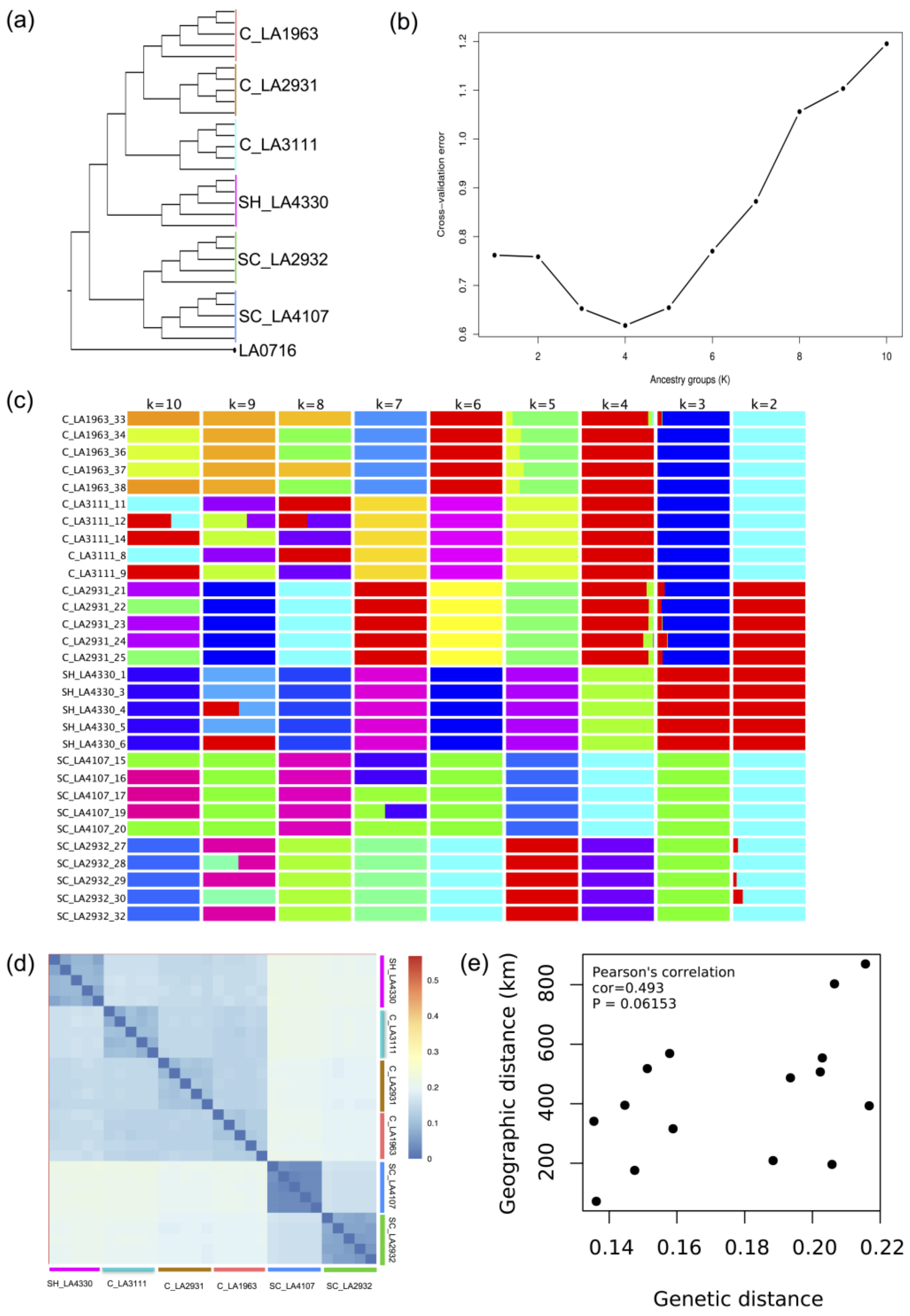


Figure S1. The relationship of different populations. (a) maximum likelihood (ML) phylogenetic tree, *S. pennellii* LA0716 is used as outgroup. (b) The accuracy of prediction for different number of co-ancestry (K). K refers to the number of presumed ancestral groups. (c) ADMIXTURE analysis showing clustering of samples from 30 individuals within K groups (2 to 10). (d) The heatmap inferring the genetic relationship using SNP data from 30 individuals. The genetic distances between pairwise individuals were calculated using Nei's distance by R package 'poppr'. (e) The correlation between genetic distance (Euclidean distance) and geographic distance.

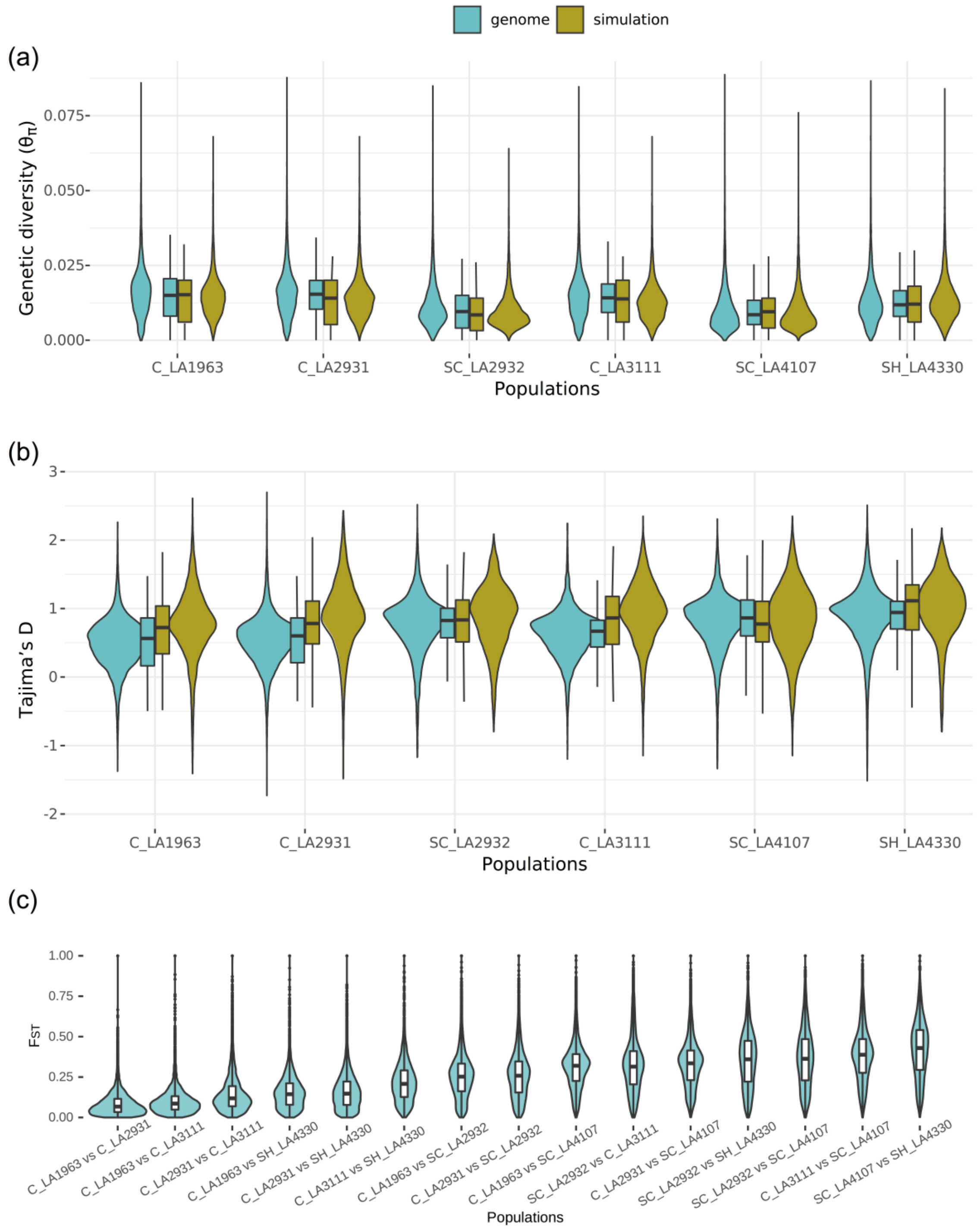


Figure S2. Summary statistics of population genetics using real SNPs and neutral simulations. (a) The distribution of nucleotide diversity ($\pi$). (b) The distribution of Tajima's D. (c) The distribution of statistics of differentiation ($F_{ST}$).

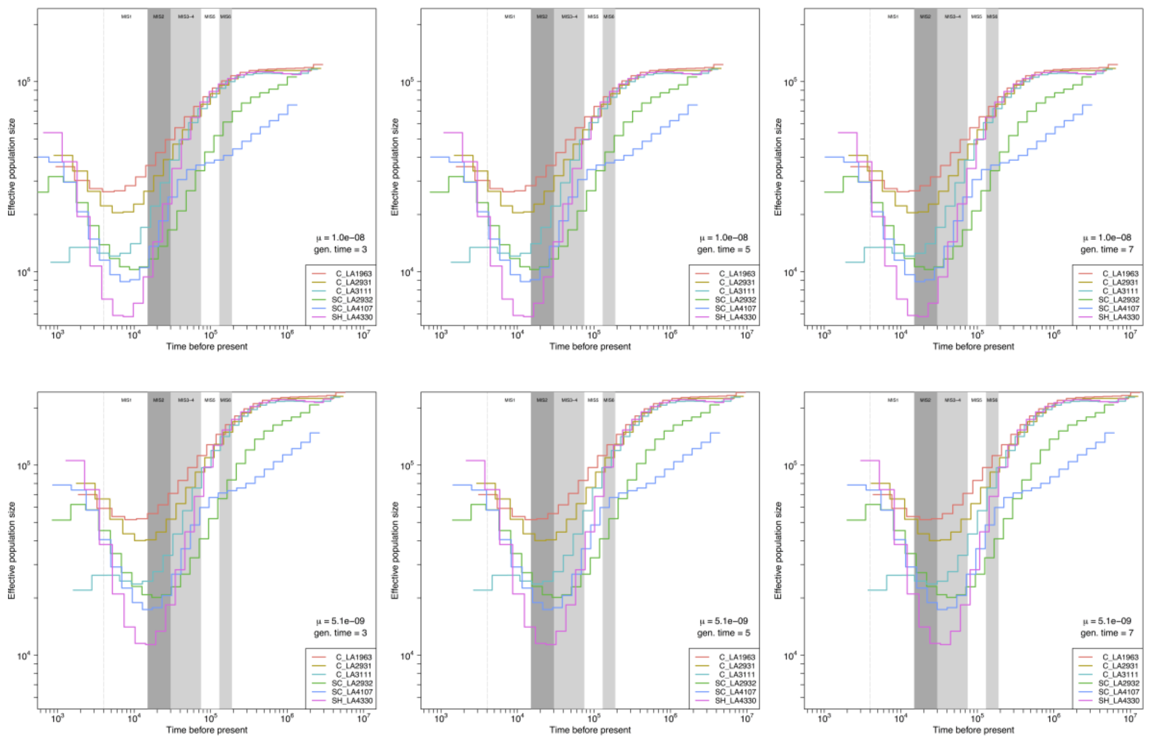

Figure S3. Estimations of effective population size through time from MSMC2 with multiple mutation rate (μ) and generation time values.

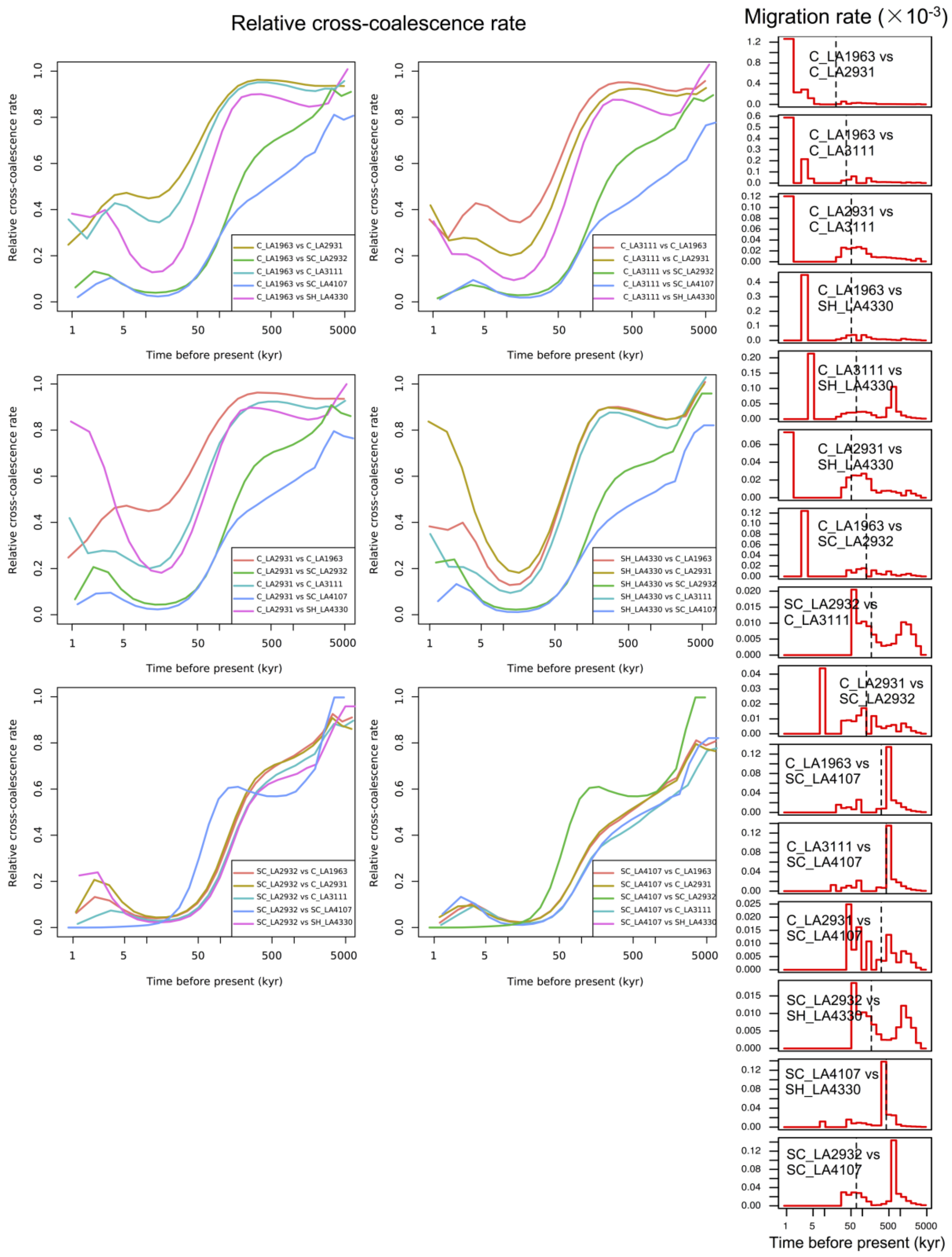


Figure S4. Relative cross coalescence rate reveals the age and pace of divergence between different populations. The two populations are well-mixed if the relative cross coalescence rate is 1 and fully separated when the value is 0. Plots in the right column indicate migration rate profiles for pairs of populations. Dashed lines indicate the time of 50% relative cross coalescence rate.

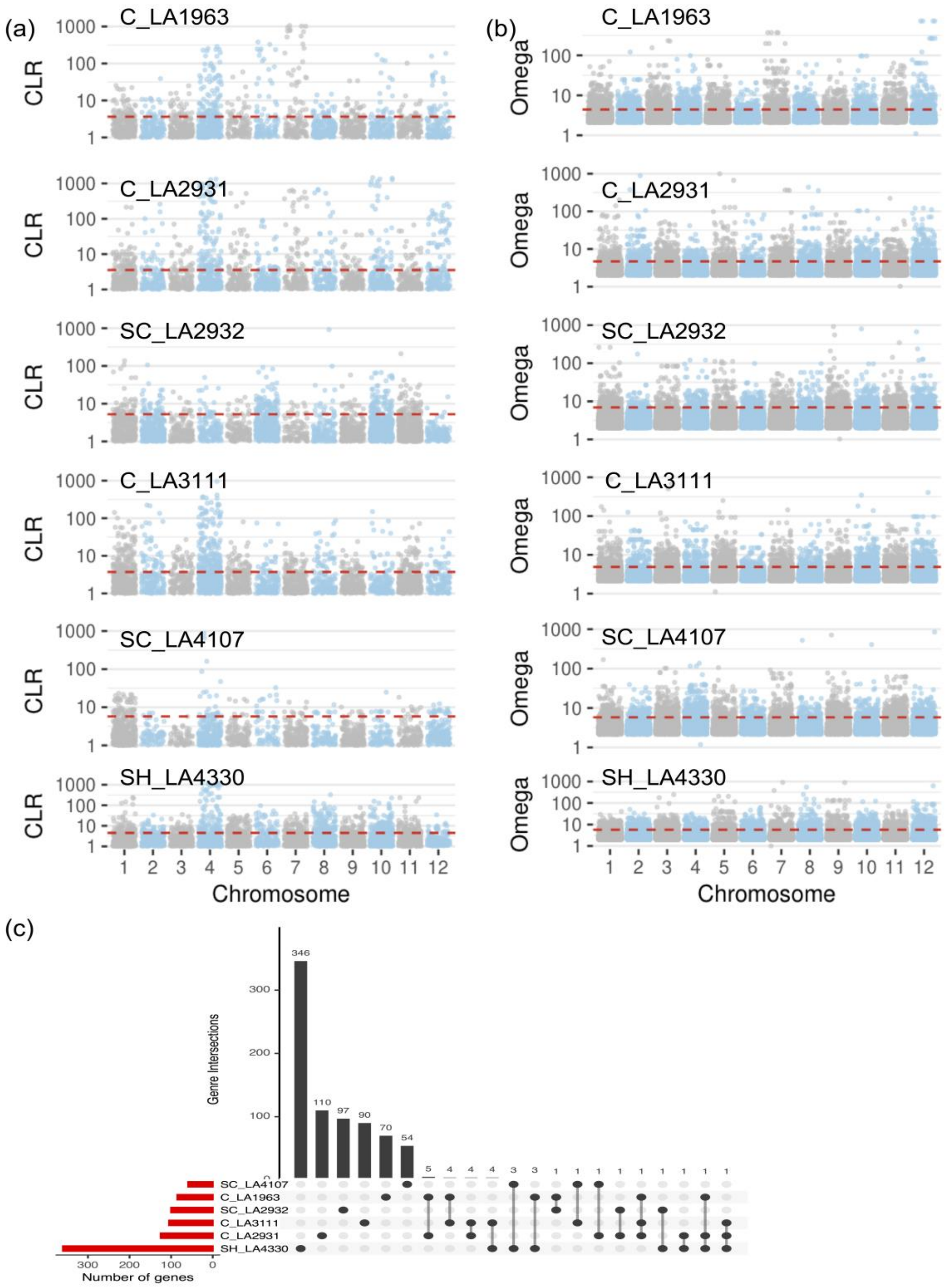


Figure S5. The plot of genome scans among six *S. chilense* populations. (a) The plot of CLR values for six populations using SweeD. (b) The plot of omega values for six populations using OmegaPlus. Dash line denotes cutoff values from neutral simulation. (c) The number of shared candidate genes between different populations. The black points represent shared genes identified in different populations.

| $2N_es$= | | 0.1 | | | 1 | | | 10 | | | 100 | | | 1000 | | |
|---|---|---|---|---|---|---|---|---|---|---|---|---|---|---|---|---|
| | | SweeD | OmegaPlus | McSwan | SweeD | OmegaPlus | McSwan | SweeD | OmegaPlus | McSwan | SweeD | OmegaPlus | McSwan | SweeD | OmegaPlus | McSwan |
| 71 kya | SH_LA4330 | 0 | 0 | 0 | 0.4 | 1.1 | 1.2 | 2.6 | 5.7 | 4.1 | 42.2 | 47.3 | 41.3 | 75.5 | 76.1 | 78.8 |
| | SC_LA4107 | 0 | 0 | 0 | 0 | 1.2 | 0.7 | 2.3 | 3.1 | 2.3 | 22.4 | 30.4 | 29.4 | 64.7 | 72.3 | 70.9 |
| | C_LA3111 | 0 | 0 | 0 | 0.9 | 0 | 0 | 1.6 | 4.4 | 2.6 | 40.8 | 41.9 | 37.8 | 70.1 | 71.8 | 73.6 |
| | SC_LA2932 | 0 | 0 | 0 | 0.4 | 0.9 | 0 | 3 | 4.9 | 3.7 | 32.4 | 43.5 | 34.7 | 73.3 | 73.8 | 72.8 |
| | C_LA2931 | 0 | 0 | 0 | 0.2 | 0.7 | 0 | 1.2 | 2.1 | 1.5 | 39.6 | 50.1 | 36.2 | 71.9 | 69.6 | 74.4 |
| | C_LA1963 | 0 | 0 | 0 | 0.8 | 0.6 | 1.1 | 2.4 | 2.3 | 1.9 | 32.8 | 46.4 | 33.7 | 67 | 67.7 | 70.5 |
| 50 kya | SH_LA4330 | 0 | 0 | 0 | 2.8 | 2.1 | 1.6 | 17.7 | 24.1 | 20.4 | 71.2 | 73.3 | 71.6 | 81.3 | 82.6 | 83.6 |
| | SC_LA4107 | 0 | 0 | 0 | 0.9 | 0.7 | 1 | 6.4 | 10.7 | 10.5 | 59.8 | 60.9 | 62.7 | 82.2 | 79.1 | 74.2 |
| | C_LA3111 | 0 | 0 | 0 | 2.2 | 1.4 | 1.4 | 14.3 | 20.5 | 13.3 | 70.9 | 72.9 | 70.4 | 76.2 | 80.7 | 77.5 |
| | SC_LA2932 | 0 | 0 | 0 | 0.7 | 0.8 | 0.7 | 8.6 | 12.4 | 10.7 | 69.4 | 67.8 | 71.5 | 82.5 | 84.1 | 80.2 |
| | C_LA2931 | 0 | 0 | 0 | 1.5 | 2.2 | 1.1 | 14.5 | 12.6 | 12.6 | 65.3 | 68.2 | 66.7 | 77.4 | 78.3 | 79.8 |
| | C_LA1963 | 0 | 0 | 0 | 0.8 | 1.4 | 1.3 | 10.4 | 15.8 | 12.5 | 67.8 | 67.3 | 69.1 | 80.4 | 79.5 | 79.3 |
| 29 kya | SH_LA4330 | 1.3 | 3.2 | 1.4 | 6.8 | 9.3 | 8.2 | 39.4 | 53.6 | 43.4 | 80.7 | 89.8 | 86.4 | 89.6 | 92.6 | 91 |
| | SC_LA4107 | 0 | 0 | 0 | 2.6 | 3.7 | 2.9 | 9.7 | 28.7 | 12.9 | 61.6 | 78.3 | 69.2 | 85.8 | 87.5 | 79.5 |
| | C_LA3111 | 0.9 | 0 | 0 | 4.4 | 6 | 5.1 | 27.7 | 49.5 | 29.3 | 75.4 | 74.9 | 81.1 | 87.3 | 90.2 | 85.1 |
| | SC_LA2932 | 0 | 0 | 0 | 3.1 | 2.4 | 3.7 | 15.3 | 34.7 | 24.2 | 75.2 | 83.1 | 77.3 | 86.4 | 88.1 | 84.3 |
| | C_LA2931 | 0 | 0 | 0 | 5 | 3.1 | 4.6 | 26.4 | 38.6 | 30.6 | 64.8 | 84.7 | 70.7 | 88.1 | 89.3 | 87.1 |
| | C_LA1963 | 0 | 0 | 0 | 2.1 | 3.4 | 2.1 | 14.8 | 29.3 | 21.3 | 69.4 | 80.5 | 71.4 | 84.3 | 92.2 | 82.2 |
| 14 kya | SH_LA4330 | 0.6 | 1.2 | 0 | 5.1 | 7.9 | 6.4 | 34.7 | 49.6 | 36.8 | 88.4 | 90.5 | 84.1 | 91.5 | 91.8 | 93.8 |
| | SC_LA4107 | 0 | 0 | 0 | 1.3 | 2.1 | 2.1 | 11.6 | 23.7 | 14.2 | 72 | 81.4 | 76.3 | 83.3 | 84.6 | 84.6 |
| | C_LA3111 | 1.1 | 0 | 0 | 3.9 | 5.4 | 4.6 | 28.3 | 40.5 | 30.6 | 83.8 | 86.5 | 86.8 | 86.7 | 93.4 | 88.2 |
| | SC_LA2932 | 0 | 0 | 0 | 1.6 | 2.8 | 1.4 | 15.6 | 30.1 | 21.3 | 76.9 | 79.2 | 80.2 | 88.6 | 87.7 | 85.3 |
| | C_LA2931 | 0.4 | 0 | 0 | 4.1 | 3.7 | 4.8 | 24.8 | 31.8 | 27.5 | 80.3 | 88.7 | 78.3 | 92.2 | 90.6 | 89.5 |
| | C_LA1963 | 0 | 0 | 0 | 1.8 | 2.9 | 2 | 16.2 | 27.3 | 20.4 | 74.1 | 79.6 | 76.6 | 89.2 | 91.4 | 90.7 |
| 8 kya | SH_LA4330 | 0 | 0 | 0 | 1.5 | 3.4 | 2.4 | 5.9 | 10.6 | 5.2 | 31.5 | 36.9 | 28.1 | 86.3 | 89.2 | 79.3 |
| | SC_LA4107 | 0 | 0 | 0 | 0 | 1.5 | 0 | 1.1 | 3.4 | 1.9 | 20.3 | 25.6 | 26.7 | 76.9 | 83.6 | 72.8 |
| | C_LA3111 | 0 | 0 | 0 | 1.3 | 3.3 | 1.6 | 2.5 | 7.3 | 4.1 | 27.6 | 32.5 | 27.8 | 81.1 | 85.4 | 76.6 |
| | SC_LA2932 | 0 | 0 | 0 | 0 | 1.1 | 0 | 1.8 | 3.6 | 2.7 | 24.4 | 34.1 | 26.6 | 79.2 | 86.3 | 76.4 |
| | C_LA2931 | 0 | 0 | 0 | 1.1 | 2.4 | 1.4 | 3.7 | 5.5 | 4.4 | 26.7 | 30.8 | 30.5 | 81.4 | 84.1 | 74.4 |
| | C_LA1963 | 0 | 0 | 0 | 0 | 1.2 | 0.5 | 2.6 | 3.9 | 2.3 | 23.5 | 27.7 | 21.4 | 78.2 | 84.2 | 72.4 |

Figure S6. The validation of our pipeline to detect selective sweeps by simulating 1,000 data sets of selective sweeps with five different selective coefficients ($N_e$ scaled) and five different ages in six populations, respectively. The numbers denote the percentage of the detected simulated sweeps using different methods.

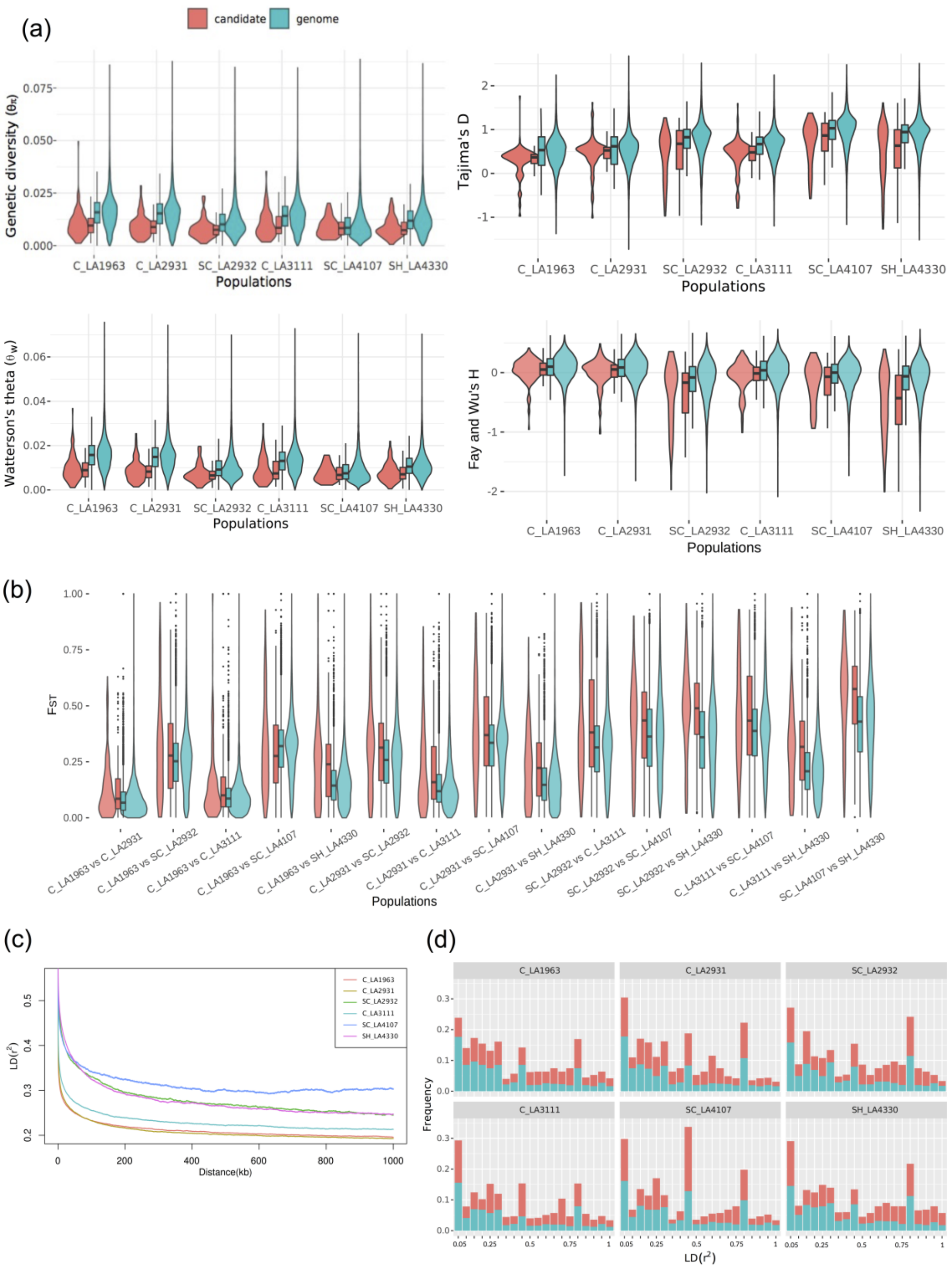


Figure S7. The comparison of statistics between whole-genome and candidate regions by 100 kb sliding windows. (a) The distribution of statistics of genetic diversity (π), Watterson's theta ($\theta_w$), Fay and Wu's H, and Tajima's D. (b) The distribution of $F_{ST}$ of whole-genome and candidate regionss. (c) LD decay against the genetic distance for pairs of linked SNP in 1000 kb distance. (d) The distribution LD of whole-genome (blue) and candidate regions (red) in 100 kb windows. Turquoise denotes whole-genome and red denotes candidate regions.

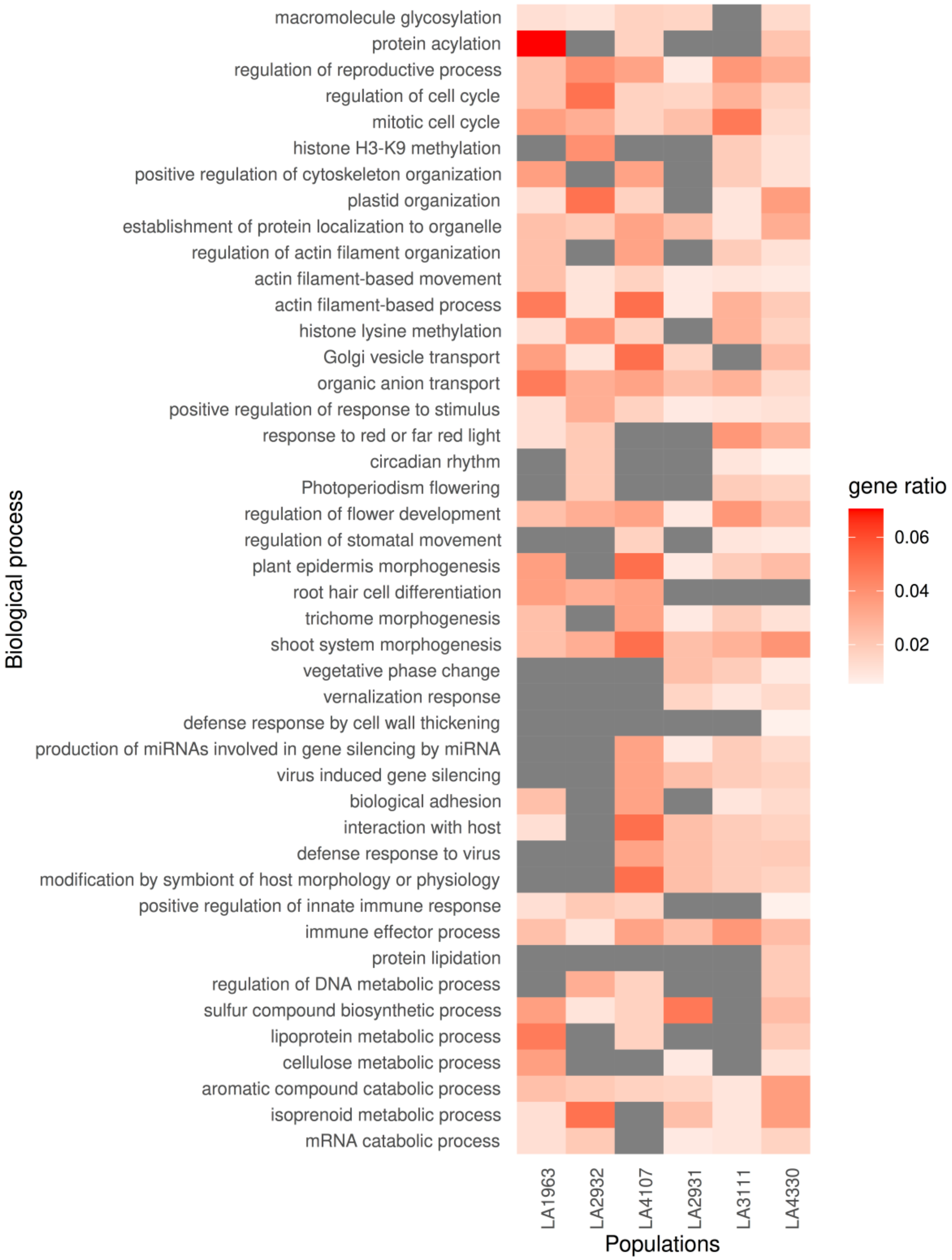


Figure S8. Gene Ontology (GO) analysis in candidate genes under positive selection enriched to the biological process in different populations. The color bar shows gene ratio in candidate genes, the gray boxes are empty and indicate that the biological process is not enriched.

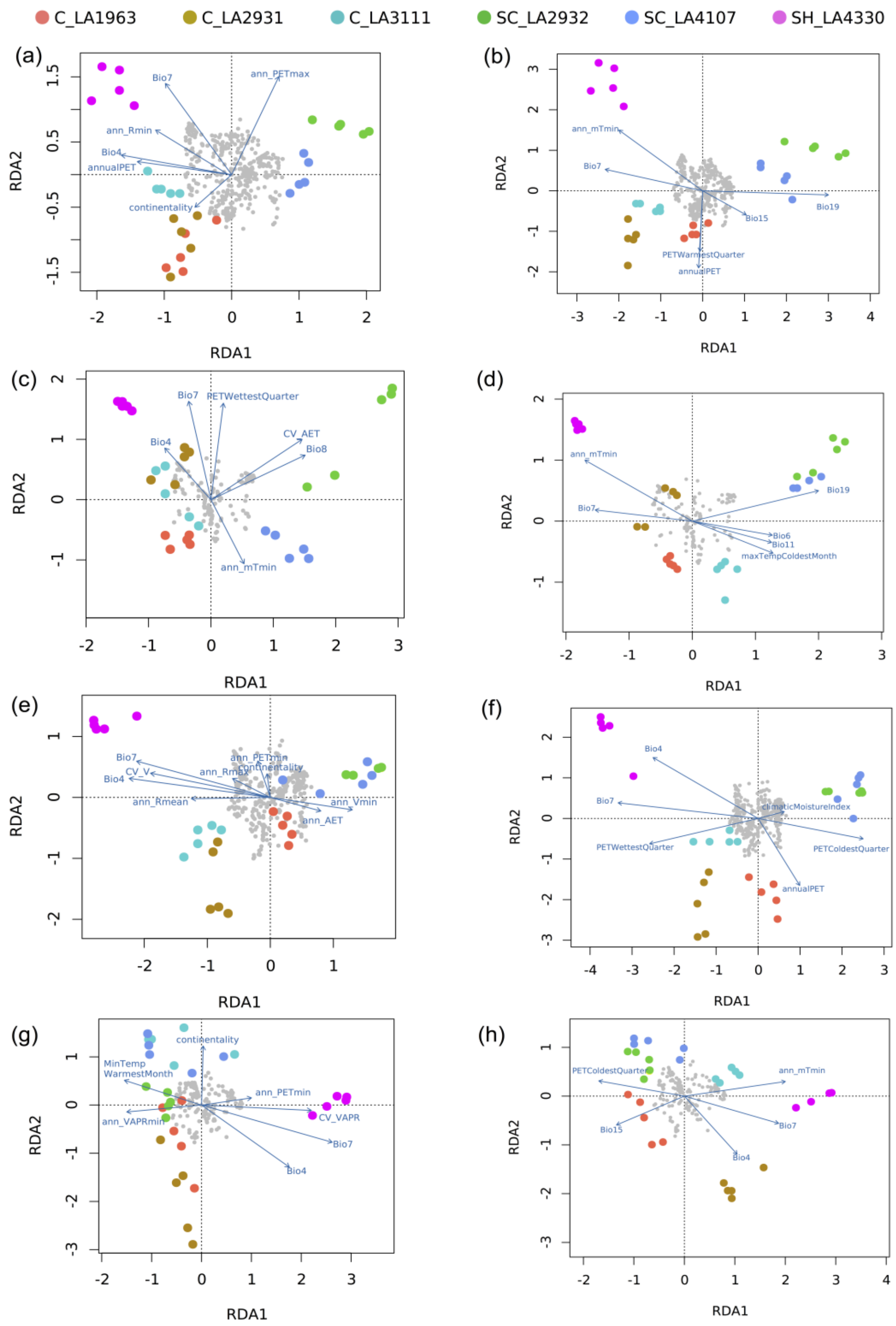


Figure S9. Redundancy analysis (RDA) of SNPs of genes related to four specific GO terms using current (a, c, e, g) and LGM (b, d, f, h) climatic variables. (a) and (b) Circadian rhythm and photoperiodism flowering. (c) and (d) vernalization response. (e) and (f) root hair cell differentiation. (g) and (h) protein lipidation. The color circles denote different populations. Arrows indicated the direction and magnitude of variables. The gray circles denote SNPs.

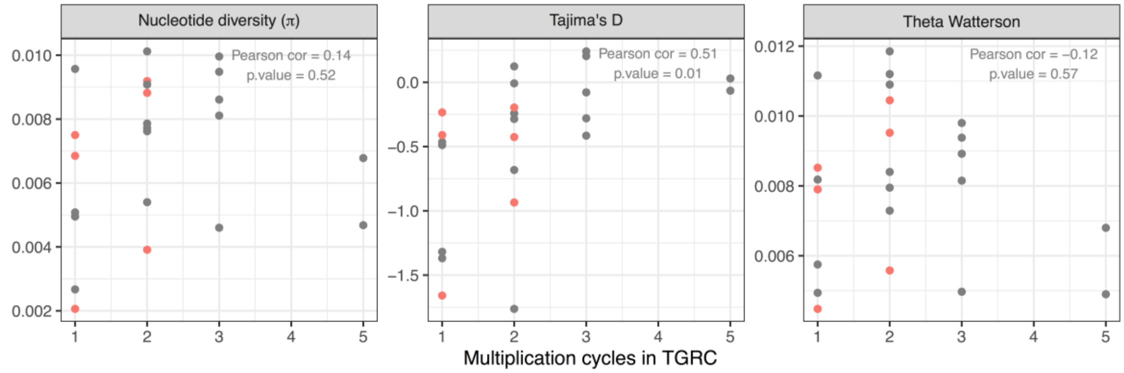


Figure S10. Population genetics statistics from the Böndel et al. (2015) for 30 sequenced loci (n=25 plants) as a function of the number of multiplication rounds at TGRC (UC Davis, USA). We thank Prof R. Chetelat for providing this information. Red points highlight the populations in common with this study.

Table S1. Population geography and habitat information

| Group | Population | Elevation (m) | Latitude | Longitude | Description | Country |
|---|---|---|---|---|---|---|
| central | C_LA1963 | 200 | -18.066667 | -70.316667 | Very dry sandy situation | Peru |
| central | C_LA3111 | 3070 | -17.466667 | -70.033333 | On side of road | Peru |
| central | C_LA2931 | 2275 | -20.916667 | -69.066667 | Dry streambed with salt or algae deposits | Chile |
| south-highland | SH_LA4330 | 3126 | -22.350833 | -68.319722 | Rocky quebrada, plants growing among rocks | Chile |
| south-coast | SC_LA2932 | 300-400 | -22.468056 | -70.225 | dry quebrada in loma zone | Chile |
| south-coast | SC_LA4107 | 86 | -25.318889 | -70.446111 | In extremely fine alluvial soil below dry waterfall | Chile |

Population information was obtained from Tomato Genetic Resource Center (TGRC) website.

Table S2. Summary statistics of variant calling in six populations

| Populations | C_LA1963 | C_LA3111 | C_LA2931 | SH_LA4330 | SC_LA2932 | SC_LA4107 |
|---|---|---|---|---|---|---|
| SNPs | 27,181,900 | 25,041,882 | 26,482,733 | 24,341,829 | 23,517,649 | 21,757,938 |
| Unique SNPs[a] | 1,162,108 | 990,787 | 1,085,306 | 9,48,174 | 1,532,616 | 1,582,572 |
| Fixed SNPs[b] | 10,245,302 | 11,371,703 | 10,513,888 | 12,130,202 | 13,096,032 | 14,707,614 |
| Insertions | 2,462,707 | 2,312,682 | 2,417,195 | 2,190,179 | 2,082,451 | 1,933,860 |
| Deletions | 2,657,687 | 2,454,456 | 2,594,730 | 2,319,887 | 2,199,316 | 2,022,334 |
| Ts/Tv[c] | 1.11 | 1.11 | 1.11 | 1.11 | 1.12 | 1.12 |
| Variant rate[d] | 30 | 33 | 31 | 34 | 35 | 38 |

[a]SNPs denote only detected in specific population.

[b]SNPs denote the conserved in all individuals of the same population.

[c]Ts/Tv denotes of ratio of Transition with Transversion.

[d]Variant rate refers to how many bases on average can detect one variant.

Table S3. The statistics of population genetics in candidate regions and whole-genome

| | nucleotide diversity ($\theta_\pi$) | | Watterson's theta ($\theta_W$) | | Tajima's D | | Fay and Wu's H | | LD ($r^2$) | |
|---|---|---|---|---|---|---|---|---|---|---|
| Population | genome[a] | candidate[b] | genome | candidate | genome | candidate | genome | candidate | genome | candidate |
| C_LA1963 | [c]$0.0165\pm8.71\times10^{-5}$ | $0.0103\pm5.35\times10^{-4}$ | $0.0164\pm7.92\times10^{-5}$ | $0.0098\pm4.98\times10^{-4}$ | $0.240\pm3.15\times10^{-3}$ | 0.163±0.029 | $0.081\pm2.21\times10^{-3}$ | 0.0348 ± 0.017 | $0.197\pm3.61\times10^{-6}$ | $0.699\pm2.43\times10^{-4}$ |
| C_LA2931 | $0.0161\pm8.71\times10^{-5}$ | $0.0096\pm4.13\times10^{-4}$ | $0.0156\pm7.88\times10^{-5}$ | $0.0088\pm3.75\times10^{-4}$ | $0.397\pm3.37\times10^{-3}$ | 0.185±0.031 | $0.066\pm2.29\times10^{-3}$ | -0.127 ± 0.018 | $0.197\pm3.85\times10^{-6}$ | $0.441\pm4.16\times10^{-4}$ |
| C_LA3111 | $0.0149\pm8.39\times10^{-5}$ | $0.0102\pm5.41\times10^{-4}$ | $0.0138\pm7.20\times10^{-5}$ | $0.0091\pm4.85\times10^{-4}$ | $0.634\pm3.47\times10^{-3}$ | 0.344±0.033 | $0.012\pm2.59\times10^{-3}$ | -0.073 ± 0.023 | $0.218\pm4.60\times10^{-6}$ | $0.663\pm4.41\times10^{-4}$ |
| SC_LA2932 | $0.0119\pm7.95\times10^{-5}$ | $0.0080\pm5.59\times10^{-4}$ | $0.0107\pm6.72\times10^{-5}$ | $0.0072\pm4.85\times10^{-4}$ | $0.770\pm4.20\times10^{-3}$ | 0.128±0.069 | $-0.134\pm3.30\times10^{-3}$ | -0.355 ± 0.062 | $0.312\pm9.19\times10^{-6}$ | $0.761\pm3.39\times10^{-4}$ |
| SC_LA4107 | $0.0105\pm7.82\times10^{-5}$ | $0.0088\pm6.20\times10^{-4}$ | $0.0090\pm6.26\times10^{-5}$ | $0.0075\pm5.15\times10^{-4}$ | $0.971\pm4.05\times10^{-3}$ | 0.530±0.064 | $-0.144\pm2.70\times10^{-3}$ | -0.172 ± 0.048 | $0.329\pm1.34\times10^{-5}$ | $0.552\pm9.33\times10^{-4}$ |
| SH_LA4330 | $0.0132\pm8.12\times10^{-5}$ | $0.0085\pm3.56\times10^{-4}$ | $0.0116\pm6.71\times10^{-5}$ | $0.0078\pm3.23\times10^{-4}$ | $0.888\pm4.25\times10^{-3}$ | 0.428±0.049 | $-0.127\pm3.50\times10^{-3}$ | -0.519 ± 0.042 | $0.316\pm7.25\times10^{-6}$ | $0.671\pm2.49\times10^{-4}$ |

[a]The values were calculated in whole-genome level, the same blow.

[b]The values were calculated in candidate regions detected by SweeD and OmegaPlus, the same blow.

[c]The values give the average and standard error of mean, the same blow.

Table S4. Pairwise $F_{ST}$ of different populations in whole genome and candidate regions

| Population | Genome | Candidate |
|---|---|---|
| C_LA1963 vs C_LA2931 | $0.058±9.48×10^{-4}$ | $0.064±9.53×10^{-3}$ |
| C_LA1963 vs C_LA3111 | $0.086±9.88×10^{-4}$ | $0.092±1.11×10^{-2}$ |
| C_LA2931 vs C_LA3111 | $0.137±1.29×10^{-3}$ | $0.165±1.23×10^{-2}$ |
| C_LA1963 vs SH_LA4330 | $0.139±1.25×10^{-3}$ | $0.196±1.21×10^{-2}$ |
| C_LA2931 vs SH_LA4330 | $0.147±1.31×10^{-3}$ | $0.172±1.10×10^{-2}$ |
| C_LA3111 vs SH_LA4330 | $0.210±1.43×10^{-3}$ | $0.267±1.27×10^{-2}$ |
| C_LA1963 vs SC_LA2932 | $0.245±1.60×10^{-3}$ | $0.258±1.70×10^{-2}$ |
| C_LA2931 vs SC_LA2932 | $0.252±1.65×10^{-3}$ | $0.288±1.61×10^{-2}$ |
| C_LA1963 vs SC_LA4107 | $0.312±1.51×10^{-3}$ | $0.390±1.59×10^{-2}$ |
| SC_LA2932 vs C_LA3111 | $0.315±1.86×10^{-3}$ | $0.367±1.91×10^{-2}$ |
| C_LA2931 vs SC_LA4107 | $0.327±1.61×10^{-3}$ | $0.351±1.75×10^{-2}$ |
| SC_LA2932 vs SH_LA4330 | $0.346±1.92×10^{-3}$ | $0.460±1.36×10^{-2}$ |
| SC_LA2932 vs SC_LA4107 | $0.362±1.94×10^{-3}$ | $0.392±2.09×10^{-2}$ |
| C_LA3111 vs SC_LA4107 | $0.384±1.79×10^{-3}$ | $0.439±1.72×10^{-2}$ |
| SC_LA4107 vs SH_LA4330 | $0.417±1.92×10^{-3}$ | $0.534±1.45×10^{-2}$ |

Table S5. The power of age estimation using sweep simulations

| Simulated age (kya) | Population | Selection strength ($2N_es$) | | | | |
|---|---|---|---|---|---|---|
| | | 0.1 | 1 | 10 | 100 | 1000 |
| 8,000 | C_LA1963 | - | 6,958±696 | 6,522±843 | 7,426±2,083 | 8522±2416 |
| 8,000 | C_LA2931 | - | 7,054±1,032 | 6,217±1,025 | 7,533±2,247 | 7486±1827 |
| 8,000 | SC_LA2932 | - | | 6,602±1,245 | 7,516±2,516 | 8936±3012 |
| 8,000 | C_LA3111 | - | 6,524±988 | 7,254±887 | 8,035±1,926 | 8,425±2,204 |
| 8,000 | SC_LA4107 | - | | 6,432±1,083 | 8,415±2,655 | 7,953±2,304 |
| 8,000 | SH_LA4330 | - | 6,162±1,124 | 7,056±1,394 | 7,756±1,994 | 7,668±1,894 |
| 14,000 | C_LA1963 | - | 11,951±1,507 | 12,458±2,216 | 16,254±2,742 | 16,307±2,983 |
| 14,000 | C_LA2931 | - | 12,540±2,041 | 12,694±2,225 | 14,358±2,851 | 16,243±3,241 |
| 14,000 | SC_LA2932 | - | 12,347±1,432 | 12,259±2,036 | 16,152±3,129 | 15,146±3,016 |
| 14,000 | C_LA3111 | - | 12,286±1,938 | 12,653±2,347 | 14,728±3,045 | 14,728±2,847 |
| 14,000 | SC_LA4107 | - | 12,166±1,486 | 12,914±2,524 | 15,433±3,635 | 16,581±3,206 |
| 14,000 | SH_LA4330 | - | 11,236±1,108 | 12,957±2,185 | 13,664±2,874 | 15,207±3,194 |
| 29,000 | C_LA1963 | - | 21,351±3,317 | 24,628±4,726 | 27,365±7,428 | 26,578±6,540 |
| 29,000 | C_LA2931 | - | 22,268±4,032 | 24,352±5,061 | 26,578±8,051 | 26,925±7,446 |
| 29,000 | SC_LA2932 | - | 23,196±3,158 | 25,325±4,328 | 29,604±8,824 | 29,460±8,527 |
| 29,000 | C_LA3111 | - | 23,147±3,640 | 24,471±4,716 | 27,368±7,413 | 26,583±7,269 |
| 29,000 | SC_LA4107 | - | 22,279±3,364 | 25,283±4,973 | 30,576±7,842 | 28,474±8,451 |
| 29,000 | SH_LA4330 | 15,736±1,726 | 20,483±3,296 | 24,365±3,724 | 27,664±7,873 | 27,916±7,435 |
| 50,000 | C_LA1963 | - | 34,589±1,207 | 42,166±3,527 | 48,251±6,640 | 48,251±6,640 |
| 50,000 | C_LA2931 | - | 36,574±1,045 | 43,796±3,425 | 47,358±7,569 | 47,358±7,569 |
| 50,000 | SC_LA2932 | - | 32,478±1,341 | 42,173±4,081 | 48,306±7,516 | 48,306±7,516 |
| 50,000 | C_LA3111 | - | 33,274±1,568 | 41,203±3,605 | 47,355±7,658 | 47,355±7,658 |
| 50,000 | SC_LA4107 | - | 37,915±1,544 | 44,379±3,842 | 52,133±9,047 | 52,133±9,047 |
| 50,000 | SH_LA4330 | - | 35,740±2,083 | 41,534±3,674 | 49,262±6,954 | 49,262±6,954 |
| 71,000 | C_LA1963 | - | 54,138±2,416 | 56,857±8,143 | 60,145±14,606 | 64,183±15,271 |
| 71,000 | C_LA2931 | - | - | 55,231±9,047 | 59,351±12,386 | 62,348±14,583 |
| 71,000 | SC_LA2932 | - | - | 56,358±8,829 | 61,434±13,296 | 66,389±16,347 |
| 71,000 | C_LA3111 | - | - | 55,406±9,451 | 58,472±13,288 | 64,182±15,592 |
| 71,000 | SC_LA4107 | - | 52,144±3,624 | 55,947±9,457 | 61,230±15,230 | 64,108±17,360 |
| 71,000 | SH_LA4330 | - | 54,641±3,347 | 53,263±10,386 | 57,263±14,,386 | 65,204±16,837 |

The numbers denote the mean sweep age ± standard deviation.

Table S6 The summary of outlier SNPs from RDA models

| RDA model | Current | | | LGM | | | Overlaps |
|---|---|---|---|---|---|---|---|
| | RDA1 | RDA2 | Total | RDA1 | RDA2 | Total | |
| All sweeps[a] | 280[b] | 3,987 | 4,267 | 350 | 1,992 | 2,342 | 1,647 |
| Circadian rhythm and photoperiod/flowering | 25 | 79 | 104 | 14 | 66 | 80 | 61 |
| Vernalization response | 7 | 19 | 26 | 0 | 23 | 23 | 18 |
| Root hair cell differentiation | 7 | 39 | 46 | 0 | 129 | 129 | 39 |
| Protein lipidation | 25 | 31 | 56 | 31 | 34 | 65 | 42 |

[a]All sweeps denote candidate regions identified overlaps between SweeD and OmegaPlus.

[b]The values denote number of outlier SNPs detected from RDA1 and RDA2.

Table S7. Summary of the number of outlier SNPs significantly correlated to climatic variables in implemented RDA

| climatic variables | All sweeps | | Circadian rhythm and photoperiodism flowering | | Vernalization | | Root hair development | | Protein lipidation | |
|---|---|---|---|---|---|---|---|---|---|---|
| | current | LGM | current | LGM | current | LGM | current | LGM | current | LGM |
| ann_AET | | | | | | | 5 | | | |
| ann_mTmin | | 249 | | 24 | 1 | 6 | | | | 18 |
| ann_PETmax | | | 13 | | | | | | | |
| ann_PETmin | | | | | | | 4 | | | |
| ann_Rmax | | | | | | | 6 | | | |
| ann_Rmean | 540 | | | | | | 4 | | | |
| ann_Rmin | | | 12 | | | | | | | |
| ann_Wmin | 510 | | | | | | | | | |
| annualPET | 1,184 | | 13 | 8 | | | | 18 | | |
| Bio15 | | | | 22 | | | | | | |
| Bio19 | 480 | 350 | | 16 | | 3 | | | | |
| Bio4 | | | 10 | | 4 | 2 | 21 | 12 | 10 | 21 |
| Bio7 | 372 | 473 | 21 | 9 | 10 | | 3 | 4 | 24 | 1 |
| Bio8 | | | | | 3 | | | | | |
| continentality | 294 | | 35 | | | | 3 | | 10 | |
| CV_AET | 386 | | | | 4 | | | | | |
| CV_R | 501 | | | | | | | | | |
| CV_VAPR | | | | | | | | | 10 | |
| MaxTempColdestMonth | | | | | | 12 | | | | |
| MinTempWarmestMonth | | | | | | | | | 2 | |
| PETColdestQuarter | | 1,090 | | | | | | 14 | | 25 |
| PETDriestQuarter | | 377 | | | | | | | | |
| PETWarmestQuarter | | | | 1 | | | | | | |
| PETWettestQuarter | | | | | 4 | | | 81 | | |
